# A TFIIB paralog drives cyclic transcription to orchestrate the archaeal cell cycle

**DOI:** 10.64898/2026.09.06.749720

**Authors:** Yin-Wei Kuo, Fabian Blombach, Frank Schult, Bettina Siebers, Finn Werner, Buzz Baum

**Affiliations:** Medical Research Council-Laboratory of Molecular Biology, Cambridge, United Kingdom; Division of Biosciences, RNAP Laboratory, Institute of Structural and Molecular Biology, University College London, London, United Kingdom; Molecular Enzyme Technology and Biochemistry, Environmental Microbiology and Biotechnology, Centre for Water and Environmental Research, Faculty of Chemistry, University of Duisburg-Essen, Essen, Germany

## Abstract

Cyclic transcription is a hallmark of the cell cycle. While transcriptional waves in eukaryotes are driven by oscillations in cyclin-dependent kinase (CDK) activity, many archaea have an ordered cell cycle despite lacking CDKs. Here in exploring how cyclic transcription is achieved in *Sulfolobus acidocaldarius,* we show that TFB2, a paralog of eukaryotic TFIIB, is cyclically expressed, associated with the promoters of division genes (including *tfb2* itself), contains a cyclin-box and is degraded as cells exit division. Furthermore, a dominant-negative TFB2 mutant interferes with division gene expression and cytokinesis. These data identify TFB2 as a central player in the circuitry orchestrating the orderly cell cycle in *Sulfolobus* and, despite the absence of CDKs, reveal parallels between the regulatory logic of archaeal cell division and eukaryotic mitosis.

## Introduction

Sequential transcriptional waves are a defining feature of ordered cell cycles, ensuring the timely expression of phase-specific genes. Coupling these waves with specific cell biological events such as genome replication, chromosome segregation and membrane remodelling is essential for high fidelity cell division. In eukaryotes, this coordination is largely achieved through the activities of CDK/cyclin complexes acting on phase-specific transcription factors such as the E2F/RB system and the MMB-FoxM1 complex in animals, reinforced by regulatory checkpoints that monitor the successful completion of key cell cycle events (*1*–*3*).

Archaea within the Thermoproteota phylum (formerly TACK archaea), which are close archaeal relatives of eukaryotes, also undergo highly ordered cell cycles (*4*, *5*). In these cells, non-overlapping phases of DNA replication and division, separated by a large G2 phase, are accompanied by cell cycle-dependent waves of transcription reminiscent of those seen in eukaryotes (*6*, *7*). However, since their genomes do not contain obvious functional homologs of CDK-cyclins, it remains unclear how these archaea regulate their cell division cycle. As a consequence, it is not yet known which if any aspects of the archaeal cell cycle control machinery were inherited by eukaryotes during eukaryogenesis.

Recent studies in Sulfolobales have identified a number of cyclically expressed archaeal-specific transcriptional inhibitors, including archaeal cell cycle regulators (aCcrs) (*8*–*10*) and the cell cycle transcription factor 1 (CCTF1) (*11*). Since these proteins arrest cell cycle progression upon overexpression, these data have been used to argue that the archaeal cell cycle is ordered by the stage-specific staggered expression of transcriptional repressors (*7*, *10*). However, this model leaves a host of questions unresolved. First, it is not known how the cyclic expression of these transcriptional regulators can ensure that key events in the cycle are complete before initiating the next phase. Second, this repressor-only model assumes the basal constitutive expression of cell cycle genes in the absence of repression, which seems surprising, and is hard to reconcile with their enrichment in the less active transcriptional compartment of the Sulfolobales genome (*10*, *12*).

Repressor-only circuits also face a deeper problem: while circuits built on negative feedback alone can, in principle, generate oscillations (*13*, *14*), robust biological oscillators resistant to noise and damping, including the eukaryotic cell cycle clock, are thought to require an interplay of positive and negative feedback loops (*15*, *16*), which drive the cycle forwards and enable irreversible transitions in cell state (*17*–*20*). Thus, fundamental questions remain: what drives periodic transcription across the archaeal cell cycle; how are transitions in cell cycle gene expression rendered irreversible and coordinated with the execution of discrete cellular events; and what does this tell us about the evolution of the eukaryotic cell cycle?

To address these questions, here we use *Sulfolobus acidocaldarius* as a model system. Our analysis identifies the cyclically expressed archaeal TFIIB homolog TFB2 as a positive transcriptional regulator of the wave of division gene expression. This study also identifies TFB2 as part of the checkpoint-like feedback mechanism that maintains division gene expression until the completion of division ring assembly. Finally, we show that TFB2 protein is rapidly degraded after passage through the ring-assembly checkpoint, triggering a change in transcriptional state, while its transcript levels decrease soon after rendering this change in cell cycle state irreversible. Together, these findings identify TFB2 as a positive master regulator of the archaeal cell-cycle regulatory network. The work also highlights features of the regulatory circuitry governing division that appear to have parallels between archaea and eukaryotes, despite the lack of obvious archaeal homologs of CDKs.

## Results

### The dynamics of cell cycle gene expression during cell division

Earlier studies examining synchronised *Sulfolobales* cultures identified sets of genes that are expressed in waves in a cell cycle-dependent manner (*6*, *7*). However, because the G1-phase in *Sulfolobales* represents only ∼5% of the cell cycle (*21*), the low levels of synchrony achievable in these experiments limited their ability to resolve gene expression during division (D-phase) from early G1 and G1/S-phase expression.

To more accurately pinpoint the timing of the expression of genes involved in cell division, we took a different approach. We took samples from a pre-synchronised population of wildtype *S. acidocaldarius* cells enriched for dividing cells, and fixed and labelled cells using antibodies against the ESCRT-III homologs (CdvB, CdvB1, CdvB2) responsible for cytokinesis (Fig. 1A). We then used fluorescence-activated cell sorting (FACS) to pool millions of these cells (∼2.5x10^7^ to 6x10^7^ cells per replicate) into five discrete phases using the initial accumulation of CdvB, CdvB1/B2 to mark the entry of D-phase, and the rapid proteasomal degradation of CdvB as an indicator of the initiation of cytokinesis (*22*). This defined five distinct states: G2-phase, early D-phase, pre-constriction, constricting and G1-phase (Fig. 1B, C).

**Figure 1.**
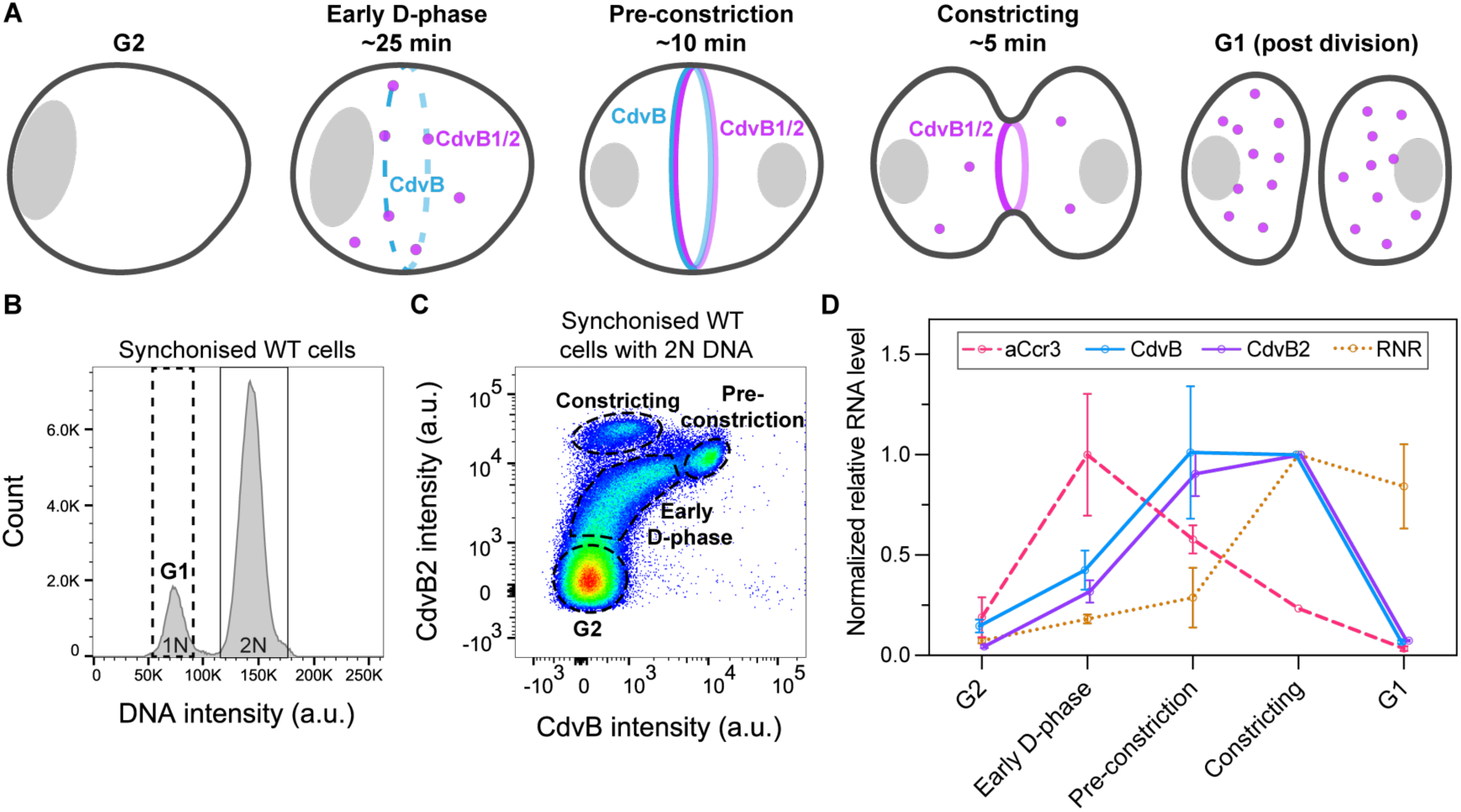
Temporal changes in cell cycle-related genes during cell division. **(A)** Schematics showing the changes in the protein levels of the ESCRT-III homologs CdvB and CdvB1/B2 during different sub-stages of cell division. Estimated time was based on cross comparing live imaging (*23*, *24*) and flow cytometry analysis of asynchronised cells. **(B)** Example flow cytometry histogram of synchronised wildtype *S. acidocaldarius* cells (DSM639) labelling the DNA content. The G1-phase cells can be separated by gating DNA staining intensity (dashed box). **(C)** Example flow cytometry scatter plot of WT synchronised cells with 2N DNA content. The sub-stages of the division phase (D-phase) can be identified by the CdvB and CdvB2 intensities. The gates indicated with dashed boxes and dashed circles were used for FACS sorting. Note G1 cells are not shown. **(D)** RT-qPCR analysis of the FACS-sorted cells showing the temporal changes of the expression of cell cycle-related genes at the RNA level. The relative abundance was normalised to the constricting phase (see methods). Error bars: mean±SDs, N=3 biological replicates.

While the RNA isolated from each of these cell populations was not sufficient for bulk RNA sequencing, we were able to perform an RT-qPCR analysis to quantify transcript levels for selected genes. This showed that while proteasomal degradation of CdvB occurs prior to the membrane constriction, CdvB and CdvB2 transcript levels peak together during cytokinesis in late D-phase. These transcripts are then rapidly lost upon completion of cytokinesis before cells enter G1-phase a few minutes later (Fig. 1D). By contrast, the transcripts for ribonucleotide reductase (RNR), which catalyses the formation of deoxyribonucleotides for DNA synthesis, reach highest levels at the D/G1-phase transition, consistent with this gene’s role in synthesizing the nucleotide building blocks required for DNA replication. Interestingly, we found that expression of the *S. acidocaldarius* homolog of the transcription factor archaeal cell cycle regulator 3 (aCcr3) (*10*) peaked prior to CdvB and CdvB2, implying an earlier role for this gene.

Taken together, these results show that cells undergo rapid changes in transcriptional state as they pass from G2 to G1 within ∼40 minutes.

### The shutoff of D-phase transcription wave requires successful assembly of an ESCRT-III division ring

Having shown that the expression of cell division genes, like CdvB and CdvB2, is rapidly switched off after completion of cytokinesis, we wondered if it would be possible to trap cells in a state with high division gene expression by arresting cells midway through the process.

Vps4 is the AAA-ATPase responsible for disassembling and remodelling ESCRT-III polymers (*25*). Previous work showed that cells overexpressing the dominant-negative Vps4 mutant (Vps4^E209Q^), despite being trapped mid-constriction, continue to progress through the cell cycle – enabling them to undergo multiple additional rounds of DNA replication (*26*, *27*). Here, in an attempt to prevent cytokinetic ESCRT-III ring assembly altogether to arrest cells earlier in the process, we overexpressed wildtype Vps4 from an arabinose-inducible promoter. As expected, this decreased the percentage of cells with division rings (Fig. 2A, B). It also arrested cell cycle progression as evidenced by the loss of G1/S-phase cells (Fig. 2C) and, in contrast to the effects of expression of the dominant negative Vps4 mutant, did not lead to a dramatic accumulation of cells with a >2N DNA content (Supplementary Fig. S1, Fig. 2C, top). These data imply that Vps4 overexpression prevents both ring assembly and cell cycle progression.

**Figure 2.**
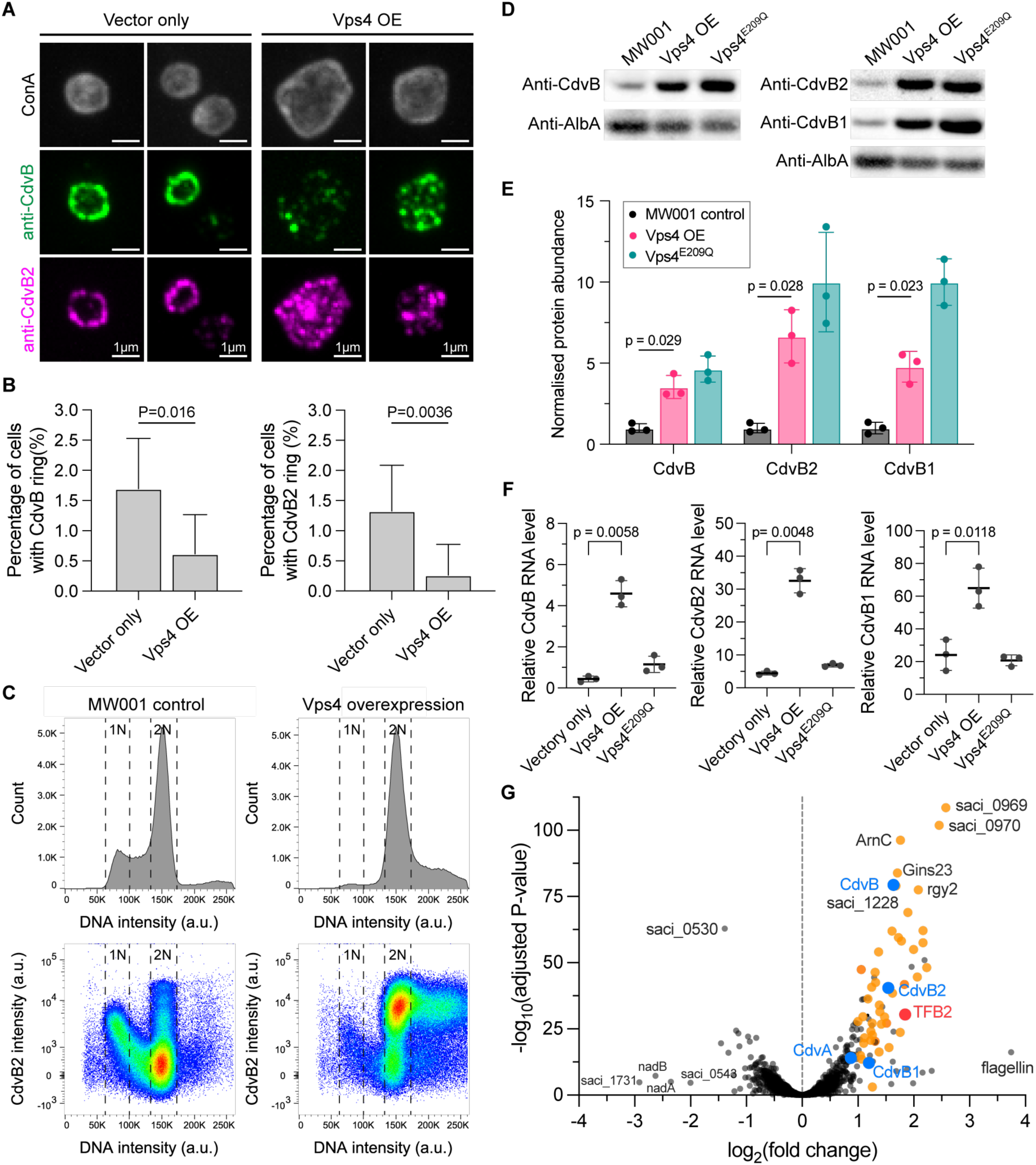
Perturbation of ESCRT-III ring assembly leads to up-regulation of division phase genes. **(A)** Example immunofluorescence images (maximum projection) of vector only control and Vps4 overexpression strain. **(B)** Quantification of cells with intact CdvB or CdvB2 rings in vector only control and the Vps4 overexpression strain (5 hr after addition of arabinose). Error bars: 95% confidence interval (CI). Fisher exact test, n= 1357 cells and 1134 cells pooled from 3 biological replicates. **(C)** Representative flow cytometry histograms and scatter plots of MW001 background strain control (left) and Vps4 overexpression strain (right, 4 hr after arabinose induction). **(D)(E)** Example Western blot of cell division-related ESCRT-III homologs (CdvB, CdvB1, CdvB2) in MW001 control, Vps4 overexpression, and Vps4 dominant negative mutant (Vps4^E209Q^), all 5 hr after addition of arabinose (D). The band intensities of the target proteins were normalised by the loading control (DNA binding protein AlbA) (E). Error bars: mean±SDs. Welch’s t-test, N=3 biological replicates. **(F)** RT-qPCR analysis of CdvB, CdvB1 and CdvB2 gene expression of vector only control, Vps4 overexpression and Vps4^E209Q^ mutant (5 hr after addition of arabinose). Error bars: mean±SDs. Welch’s t-test, N=3 biological replicates. **(G)** Volcano plot showing differential gene expression based on an RNA-seq analysis that compared the impact of Vps4 overexpression relative to Vps4^E209Q^ mutant expression after 5 hr of arabinose induction (N=4 biological replicates each). The highly up-regulated (fold change ≥ 2) cyclic genes were labelled in orange, the cell division genes (CdvA, CdvB, CdvB1, CdvB2) were highlighted in blue, and TFB2 was shown in red.

In exploring the cause of the cell cycle arrest induced by Vps4 overexpression, we discovered that although the depolymerisation of ESCRT-III division rings might be expected to trigger monomer degradation (*11*, *26*), when analysed by flow cytometry, CdvB2 protein was seen accumulating to high levels in Vps4 OE cells – equivalent to those seen in unperturbed cells entering late D-phase (Fig. 2C, bottom). The same result was revealed by Western blotting where, following the over-expression of Vps4, the full set of ESCRT-III homologs (CdvB, CdvB1, CdvB2) accumulated to levels similar to those seen in cells expressing the hydrolysis-deficient dominant negative Vps4^E209Q^ mutant (Fig. 2D, E).

The cause of protein accumulation was different in the two cases. An RT-qPCR analysis showed that Vps4 OE was associated with a drastic up-regulation in the levels of transcripts for all three ESCRT-III homologs (Fig. 2F), as expected if cells were trapped mid-division with high levels of division gene expression. By contrast, Vps4^E209Q^ expression prevented proteasome-mediated degradation of ESCRT-III monomeric proteins (*26*), keeping protein levels high even though division gene transcription was switched off as cells progress into G1 and S-phase (Fig. 2F).

To test whether the ability of Vps4 overexpression to arrest cells mid-division was due to its impact on division ring formation, we explored the impact of blocking ring formation in different ways using constructs that interfere with the interaction between the CdvA template ring and CdvB – either by expressing a dominant negative CdvA mutant (CdvAΔE3B) that lacks the C-terminal CdvB binding site (E3B motif) (*24*, *28*) or by expressing β-galactosidase (LacS) fused to the C-terminal region of CdvB (LacS-CdvB^194-end^) which competitively inhibits interactions between endogenous CdvB and CdvA (Fig. S2A) (*11*). When immunofluorescently-labelled cells were imaged, we were able to confirm that the two constructs interfere with CdvB and CdvB2 ring formation (Fig. S2B, C). This was despite the fact that the expression of either CdvAΔE3B or LacS-CdvB^194-end^ led to an increase in the levels of CdvB and CdvB2 transcripts along with the corresponding proteins (Fig. S2D-G), mirroring the impact of Vps4 overexpression. These results suggest that, while cells with a fully assembled division ring that is unable to constrict can enter a G1/S phase transcriptional state, the failure to complete division ring assembly locks cells in the Division phase.

To better understand the nature of the mid-division arrest, we used RNA-Seq to compare the transcriptomes of cells over-expressing wildtype Vps4 versus Vps4^E209Q^. While these two proteins differ at a single amino acid, this has profound consequences for gene expression. As observed by RT-qPCR, Cdv gene expression is up-regulated following Vps4 OE strain, in a way that is not seen in the Vps4^E209Q^ strain (Fig. 2G, blue dots). Additionally, the set of highly up-regulated genes (log_2_(fold change) ≥ 1) observed following Vps4 OE largely overlapped with the set of cyclically expressed genes previously found to peak around D-phase (*6*, *29*), and with the set of genes annotated in arCOG database (*30*) to have cell cycle and DNA replication related functions (48 out of 75 genes; Fig. 2G, orange dots). This includes the protein kinase ArnC (*31*, *32*), the TFIIB family protein TFB2, the DNA replication complex protein Gins23 (*33*, *34*), the OLD family ATPase Cran1 (*35*), dCTP deaminases, the chromatin protein Sul12a (*36*), and the chromosome dimer resolution recombinase XerA (*37*) (Supplementary data S1). Notably, amongst the top upregulated genes were the cyclically expressed conserved operon saci_0969/0970, whose functions remain to be determined.

Taken together, these results suggest that a failure to assemble a functional cytokinetic ring in *Sulfolobus* triggers a checkpoint-like response that maintains the D-phase transcription wave in the ON-state in a way that aids ring completion. As previously suggested based on an analysis of DNA segregation (*24*), this means that the completion of division ring assembly functions as a critical regulatory step during cell division: it is a prerequisite for switching off division genes expression to allow for D-phase exit and to prepare for entry into G1 and S-phase.

### Discovery of TFB2 as a cell cycle regulator

Having found a way to lock *Sulfolobus* cells in a cell cycle phase in which division gene expression is active, we searched for potential positive regulators of the transcriptional wave amongst the upregulated genes. This identified the basal transcription factor TFB paralog, TFB2, as a potential regulator of transcription that is expressed at relatively high levels in the arrest (Fig. 2G, Fig. S3A-C).

Archaeal TFB factors are homologs of the eukaryotic TFIIB that, together with the TATA-binding protein (TBP), are necessary and sufficient for promoter recognition and start site-specific transcription initiation *in vitro* (*38*, *39*). Many archaea encode several TFB variants that regulate distinct subsets of genes, albeit with a degree of redundancy (*40*). TFB2 is a TFB paralog in *Sulfolobus* whose expression levels were previously found to be cyclically modulated at mRNA level (*6*). To better understand the evolutionary relationships of TFB2 homologs across archaea and eukaryotes, we performed a phylogenetic analysis (Fig. S4). In agreement with the analysis carried out by Koonin and colleagues (*41*), this implies that TFB2 arose via a gene duplication early on in the evolution of TACK archaea (Fig. S4), many of which have ordered cell cycles (*6*, *21*, *42*).

In order to test whether TFB2 is associated with promoters of cell division genes, under conditions of high TFB2 expression levels, we carried out ChIP-seq using a StrepII-tagged TFB2 variant in cells overexpressing Vps4, and compared it to the binding profiles of TFB1, which is responsible for the transcription of most housekeeping genes in *Sulfolobales* (*43*). TFB2 was specifically enriched at promoters of genes of key cell division machinery including the *cdvABC* operon and the ESCRT-III homologs *cdvB1 and cdvB2* (Fig. 3A, S5A). At genome-scale, TFB2 peaks were strongly enriched in the promoter region of genes whose expression is upregulated following Vps4 overexpression thus connecting TFB2 binding and transcription output (Fig. 3D, S5B). Unbiased peak calling revealed only limited overlap with promoters bound by TFB1 (Fig. S5C), and the occupancy of TFB1 and TFB2 was not strongly correlated (Fig. 3Β).

**Figure 3.**
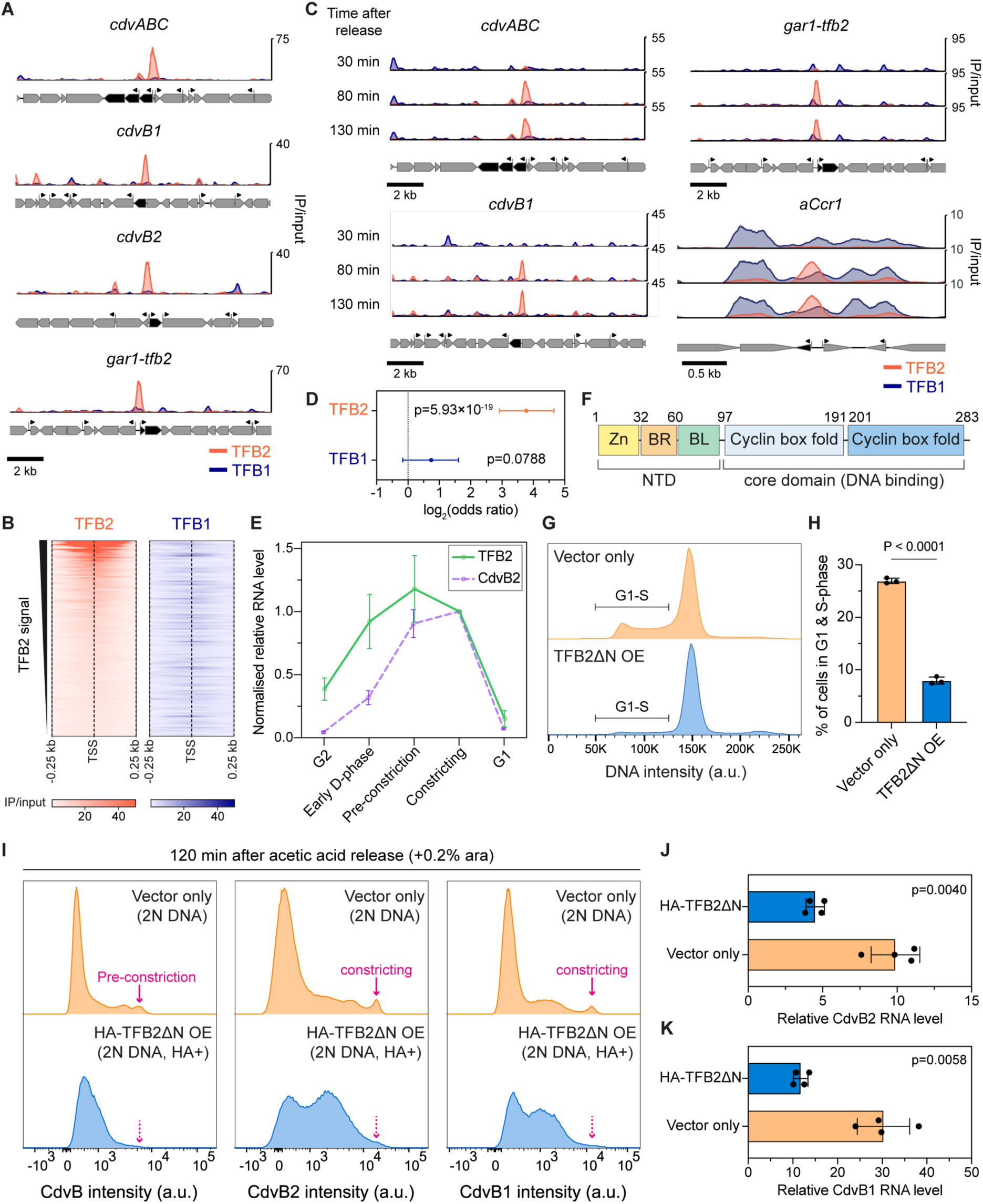
TFB2 regulates expression of cell division genes. **(A)** TFB2 (orange) and TFB1 (blue) occupancy at the promoters of cell division genes in Vps4 overexpression cells. Signal represents the mean of at least two biological replicates. **(B)** Heatmap of TFB2 and TFB1 ChIP-seq occupancy at TFB2-occupied promoters. Representative heatmaps from one biological replicate for 123 promoters are shown. For TFB peaks overlapping with pairs of closely spaced divergent promoters the promoter with higher expression level is shown. TSS: transcription start site. **(C)** TFB2 (orange) and TFB1 (blue) occupancy at the promoters of selected cyclically expressed genes in synchronised cells at T= 30, 80 and 130 min after released from acetic acid arrest. **(D)** Association of TFBs occupancy in Vps4 OE cells with upregulated operons in Vps4 OE relative Vps4^E209Q^ mutant cells (see Methods for details). Fisher’s exact test; error bars represent 95% confidence intervals determined by Fisher’s method. **(E)** RT-qPCR analysis of sorted cells (see Fig. 1) showing changes in TFB2 transcript abundance as cells progress through the division phase. Note that the CdvB2 curve from Fig. 1D was plotted again here for comparison. Error bars: mean±SDs. N=3 biological replicates. **(F)** Domain architecture of *S. acidocaldarius* TFB2 (*45*). **(G)** Example flow cytometry histograms of vector only control and cell expressing TFB2ΔN (3 hr after addition of arabinose induction). **(H)** Quantification of the proportion G1-S phase cells (gates shown in (G)). Error bars: mean±SDs. Welch’s t-test, N=3 biological replicates each. **(I)** Example flow cytometry histograms of synchronised vector only control strain, and strain expressing TFB2ΔN with an N-terminal HA-tag with 2N DNA content. Arabinose induction was started 2 hr before release from acetic acid arrest. Samples collected at around the peak of division (T=120 min after release) were shown here. For the TFB2ΔN mutant, HA-positive cells were gated to account for expression heterogeneity. **(J)(K)** RT-qPCR analyses of synchronised vector only control and HA-TFB2ΔN expressing cells at around the peak of division. Note that one time point earlier than (I) was used to account for the lag between transcription and protein expression. Error bars: mean±SDs. Welch’s t-test, N=4 biological replicates.

To characterise TFB2 recruitment under physiologically relevant conditions in cycling cells, we performed ChIP-seq on synchronised populations of cells at different time points (T=30 min, 80 min and 130 min) following release from a G2 arrest. This analysis showed that cyclically expressed genes, including most previously defined cell division genes, exhibit an increase in TFB2 binding around the peak of division (Fig. 3C, Fig. S5D, T=80 and 130 min), indicating that promoter binding is correlated with activation of the D-phase transcription wave. In addition to the expected targets of division gene expression, the TFB2 regulon also identified several cyclically expressed genes of unknown function (Supplementary data S2), increasing the set of cell cycle-related machinery that have yet to be characterised. Using the MEME algorithm, we sought to identify distinctions between DNA sequence motifs. As expected, the TBP-binding TATA boxes were near-identical in TFB2- and TFB1-responsive promoters. The B-recognition elements (BRE) also appeared similar, but with a different spacing between these two motifs suggesting distinct promoter architectures (Fig. S5E).

Having identified the *tfb2* gene itself as a target of the division wave (Fig. 3A, C), to determine more precisely the timing of TFB2 expression during a normal cycle, we carried out an RT-qPCR analysis of sorted synchronised wild type cells. This revealed that TFB2 transcripts accumulate in parallel with CdvB and CdvB2, peaking in late D-phase, before rapidly falling as cells enter G1 (Fig. 3E), in line with earlier data showing that TFB2 is part of the division gene wave (*6*). Next we wanted to determine if TFB2 protein levels reflect the levels of RNA or if TFB2 is also subject to cyclic degradation, as was suggested previously (*11*, *22*, *32*). The endogenous TFB2 levels were too low to be detected via flow cytometry, and to overcome this challenge we overexpressed a StrepII-tagged TFB2 variant from a plasmid that drives flat and inducible expression (*11*). This did not disrupt cell division or cell cycle progression (Fig. S3D), and strikingly, StrepII-tagged TFB2 levels decreased sharply around the pre-constriction phase of division (Fig. S3E-G), at a similar time to CdvB. These data demonstrate that TFB2 undergoes rapid protein degradation as cells complete the division process. The rapid degradation of TFB2 protein during division was dependent on the protein’s C-terminal cyclin fold, since the level of a truncated mutant lacking this portion of the protein (TFB2ΔC) remained high throughout D-phase in comparison to the full length TFB2 (Fig. S3H-J). The fact that cyclin folds are targeted for cyclic degradation in archaea (TFB) and eukaryotes (cyclins) hints at a recurring evolutionary strategy of cell cycle regulation.

Taken together, these data show that *Sulfolobus* cells entering D-phase accumulate both TFB2 transcript and protein. Through TFB2 binding to its own promoter (part of the *gar1-tfb2* operon (Fig. 3C)), this likely triggers a positive feedback loop that advances cells into division. Following division ring assembly, TFB2 protein is then rapidly degraded by the proteasome alongside CdvB as cells undergo cytokinesis, in a manner that depends on the presence of a cyclin fold, while TFB2 mRNA levels remain high. Finally, TFB2 and CdvB transcripts levels are then rapidly lost as cells enter G1 prior to entry into S-phase.

As *tfb2* is an essential gene (*44*), we could not obtain a deletion mutant to explore the loss of function phenotype. As an alternative approach, we constructed a dominant negative version of the protein, TFB2ΔN, which is N-terminally truncated and lacks the ability of recruit RNA polymerase but is able to engage with promoter binding (Fig. 3F). Flow cytometry analysis showed that the induction of TFB2ΔN expression strongly disrupted cell cycle progression, leading to a large reduction in G1-S-phase cells (Fig. 3G, H), as would be predicted if TFB2 is a positive regulator of division gene expression. Furthermore, cells expressing an HA-tagged version of the TFB2ΔN mutant protein that entered D-phase, were unable to express high levels of CdvB/B1/B2 to complete division (Fig. 3I, S6A,C).

Consistent with the role for TFB2 in the induction of division gene expression, an RT-qPCR analysis revealed that transcript levels of division genes CdvB1 and CdvB2 were significantly reduced relative to their expected peak expression in cells expressing the TFB2ΔN mutant during both a synchronous cycle (Fig. 3J, K). Since levels of HA-TFB2ΔN were variable in the population in ways that could interfere with this analysis, we also compared the levels of CdvB1/B2 protein abundance with that of TFB2ΔN in an asynchronous population. This revealed a negative correlation in the expression level of TFB2ΔN and CdvB1/B2 proteins (Fig. S6B), consistent with the truncation acting as a competitive inhibitor of TFB2-dependent division gene expression. This conclusion was further supported by immunofluorescence microscopy, which showed that cells expressing the TFB2ΔN mutant were unable to assemble full division rings (Fig. S5D). Taken together, these results suggest that TFB2 likely drives the expression of division genes necessary for successful cell division in a way that can be competed out by the expression of a dominant negative construct.

### The transcription factor aCcr1 shuts off TFB2 target gene expression and promotes transition to G1/S-phase

Having shown that TFB2 levels rise as cells enter division, inducing expression of the TFB2 regulon, including TFB2 itself, and that TFB2 is subsequently degraded following ring assembly, we wanted to better understand how the D-phase transcription wave is switched off as cells enter G1. We focused this analysis on aCcr1, which is part of the TFB2 regulon (Fig. 3C) and whose mRNA levels were reduced by TFB2ΔN overexpression (Fig. S7A). This made aCcr1 a candidate regulator of the D-phase transcriptional wave, as previous work has identified it as a transcriptional repressor that can block the expression of *cdvA* and other genes when over-expressed in *Sa. islandicus* (*8*, *9*) and *S. acidocaldarius* (Fig. S7B). To assess the role of aCcr1 in exit from D-phase, we first characterised its temporal expression profile using RT-qPCR across an unperturbed cell cycle. In FACS sorted wildtype cells, aCcr1 mRNA levels were found to peak in late D-phase. Importantly, these data suggest that this repressor accumulates shortly after TFB2 (Fig. S7C). This suggested the possibility that the expression of aCcr1 might serve to break the TFB2 positive feedback loop and prepare cells for exit from division. This would be aided by the early degradation of TFB2, which precedes loss of TFB2 transcript.

We then examined the impact of aCcr1 overexpression which led to the rapid loss of G1/S-phase cells, followed by accumulation of cells with more than 2N DNA content (Fig. 4A, B, Fig. S7D), indicating that aCcr1 blocks cell division while allowing for continuing DNA replication and relicensing. Furthermore, cells over-expressing aCcr1 were much larger in size than the vector only control (Fig. S7E), consistent with them undergoing continued growth despite failing cell division. A transcriptomic analysis of this experiment further showed that many of the D-phase genes which are part of the TFB2 regulon, including *tfb2* itself, were downregulated in aCcr1 overexpressing cells (Fig. 4C, blue). By contrast, and consistent with the expression phenotype of aCcr1 overexpression, genes associated with S-phase entry, including RNR, replication origin binding protein UBP (saci_0847) (*46*), and DNA repair proteins Nre (saci_1142) and RadA (saci_0715), (*6*) were upregulated following aCcr1 overexpression (Fig. 4C, red). These data support the idea that aCcr1 promotes the exit from division phase and entry into G1.

**Figure 4.**
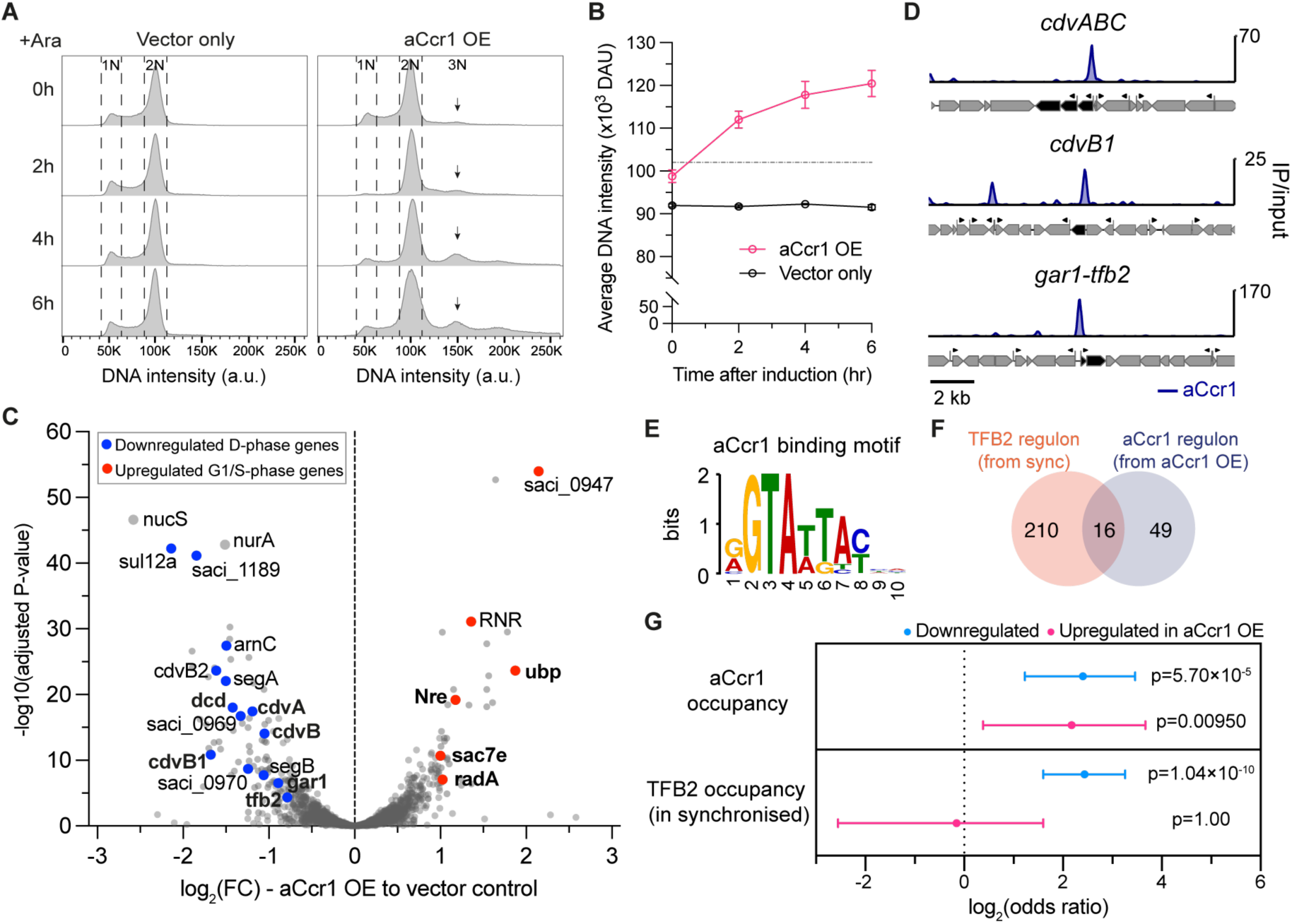
aCcr1 represses TFB2-dependent division gene expression and promotes transition into G1/S-phase. **(A)** Example flow cytometry histograms of vector only control and aCcr1 overexpression during the time course of arabinose induction. **(B)** Quantification of average DNA intensity of strains shown in (A). Error bars: mean±SDs, N=3 biological replicates. **(C)** A volcano plot showing differential gene expression of aCcr1 OE relative to empty vector control based on RNA-Seq experiments (Supplementary data S1). Selected up- and down-regulated genes associated with cell cycle function are marked in red and blue, respectively. Genes with aCcr1 occupancy at the promoter region (e.g. CdvA) are shown in bold. All RNA-Seq samples were from 3 hr after arabinose induction. N= 4 biological replicates each. **(D)** Example aCcr1 occupancy at the promoter of division genes in aCcr1 OE cells. **(E)** aCcr1 binding motif identified from ChIP-seq experiment using MEME (*48*). **(F)** Venn diagram of TFB2 and aCcr1 regulons from the peak calling data of ChIP-seq experiments (Supplementary data S2). TFB2 regulon is based on peak calling at T80 and T130 in the synchronised cells (Fig. 3C), and the aCcr1 regulon is determined by ChIP-seq experiment of aCcr1 OE cells. **(G)** Association of aCcr1 occupancy (in aCcr1 OE) and TFB2 occupancy (in synchronised cells at 80 min after release) at the promoter regions with highly differentially regulated operons in aCcr1 OE relative to vector only control (see Methods for analysis details). Error bars and p-values: 95% CI with Fisher’s exact test.

To gain insight into the underlying occupancy of aCcr1 at promoters, we performed ChIP-seq for the aCcr1 overexpression strains using polyclonal aCcr1 antibodies. ChIP-seq data for aCcr1 showed clear binding at the promoter regions of division genes including *cdvA, cdvB1,* and *gar1-tfb2* (Figure 4D, Supplementary data S2), with a binding motif (Fig. 4E) near identical to the previously identified *Sa. islandicus* aCcr1-box (*9*) and *S. acidocaldarius* CCR-2 box (*47*). In many cases, including *cdvA* and *gar1-tfb2* (Figure 4D), two to three aCcr1-boxes were identified at the same promoter (Fig. S7G). While aCcr1 peaks were found in the promoter regions of several D-phase genes (Fig. 4D), aCcr1 and TFB2 regulons showed only a small overlap (Fig. 4F; examples in Fig. 4C, bold). Consistent with its suggested role as a transcriptional repressor, there was a significant association of aCcr1 with the promoters of highly downregulated genes in aCcr1 OE cells (Fig. 4G, 14 out of 65 genes with fold change ≤ 0.5), including the DNA repair nucleases *nurA* (*saci_0148*) and *nucS* (*saci_0200*) (Fig. 4C, S7F). Importantly, promoter binding of TFB2 in the synchronised cells was also associated with highly downregulated genes in aCcr1 OE cells (Fig. 4G), encompassing half of the genes with fold-change ≤ 0.5 (32 out of 65 genes). These results indicate that aCcr1 directly suppresses a small subset of D-phase genes including *tfb2* itself. These data help to show how the D-phase transcription wave is switched-off by aCcr1 following the downregulation of *tfb2* expression and TFB2 degradation. Interestingly, aCcr1 promoter occupancy was also associated with a small number of upregulated genes (Fig. 4G, 5 out of 29 genes with fold change ≥ 2). Four of these genes are linked to DNA replication, repair or found to be cyclically expressed around S-phase (Fig. 4C bold, Fig. S7H) (*6*), suggesting that aCcr1 may also play a role, in a way that remains to be elucidated, in activating S-phase genes in addition to repression of D-phase gene expression.

These data support a model where the accumulation of TFB2 protein in early D-phase triggers the expression of division ring proteins along with aCcr1. Division ring formation and passage through the ring assembly checkpoint then trigger TFB2 degradation, initiating exit from the division program. At this point in the division process the presence of aCcr1 acts as a delayed negative feedback circuit to repress *tfb2,* breaking the positive feedback in the system. At the same time, it shuts down the expression of a small set of important division genes to help drive exit from division program and to promote entry into G1 and the initiation of S-phase. Together, these two regulatory steps ensure timely silencing of division genes while triggering progression into the next round of the cell cycle.

## Discussion

Our results identify TFB2 as a major regulator of division phase (D-phase) transcription in *Sulfolobus acidocaldarius* and reveal a checkpoint-like mechanism coupling the D-phase transcriptional program to the key event of cell division. We show that TFB2 is expressed during D-phase (Fig. 3E) rather than at the G1/S transition as previously proposed (*6*). TFB2 remains highly expressed in division ring assembly mutants (Fig. 2G, S3A-C), accompanying sustained upregulation of D-phase genes (Fig. 2G). Using a ChIP-seq analysis we have also defined the TFB2 gene regulon. This includes promoters of cyclically expressed genes (Fig. 3A, C), and includes *tfb2* itself. Finally, expression of a dominant-negative TFB2 mutant demonstrated that TFB2 is required for full activation of D-phase genes, including *cdvB1* and *cdvB2* (Fig. 3I–K). Collectively, these findings establish TFB2 as a positive regulator of D-phase transcription.

The identification of TFB2 as an archaeal cell cycle regulator raises the question of how it acquired this role during evolution. Archaeal transcription is thought to have evolved from a simple system comprising TATA-box binding protein (TBP), a single TFB/TFIIB, and TFE/TFIIE (*38*). In many archaea, duplication of *tfb* genes enables functional specialization of its paralogs (*49*, *50*). This process of gene duplication and divergence continued in eukaryotes, and accompanied the diversification of RNA polymerases (*38*). In the case of Thermoproteota (formerly the TACK archaea), TFB2 retains its conserved syntenic association with *gar1*, a gene involved in the biogenesis of ribosomes, suggesting that *Sulfolobus* TFB2 represents the ancestral copy and experienced accelerated evolution after paralogization (*41*). These observations are consistent with an evolutionary pathway in which duplication of TFIIB enabled one paralog (TFB2) to acquire specialized cell cycle function while the duplicated paralog (TFB1) retained its ancestral role in basal transcription. While the mechanistic rationale underlying this specialisation awaits high resolution structures of the TFB1 and TFB2 pre-initiation complexes, divergence of their shared cyclin box-containing domain may play a key role in their function. As shown in this study, this cyclin-homology domain plays not only a key role for basic transcriptional function via DNA binding, but similar to eukaryotic cyclins (*51*, *52*), facilitates the proteasome-dependent degradation of TFB2 during exit from division phase. This regulation is likely specific to the TFB2 cyclin domain as TFB1 is not subject to similar cyclic degradation.

Intriguingly, several central cell-cycle regulators, including cyclins and RB-family proteins, contain cyclin-box folds that is most likely derived from the ancient TFB/TFIIB fold (*53*–*55*). This makes archaeal TFB proteins the most probable ancestors of cyclins. In eukaryotes, cyclin-box-containing proteins can also serve as central components of the cell cycle transcriptional programs though primarily through their association with CDKs which phosphorylate both transcription factors and RNA polymerase II (*56*–*58*). While a detailed analysis of similar proteins across the archaea and in Asgard archaea will likely be required to resolve the evolutionary path of the cyclin fold, it is striking to see the parallels between the functional specialization leading to the cyclic expression of TFB2, which plays a major role in archaeal cell cycle control, and the emergence of cyclin–CDK regulation in eukaryotes.

This study’s identification of TFB2 as the general transcription factor driving cyclic division gene expression in *Sulfolobus* reveals several important regulatory features of the archaeal cell cycle. First, the D-phase transcription wave is rapidly switched off following the completion of cell division within ∼5% of cell cycle time (Fig. 1D), indicating tight temporal coupling between cytokinesis and transcriptional inactivation. Second, the phenotype of the division ring assembly mutants (Fig. 2, Fig. S2) and the Vps4 dominant negative mutant (*26*) indicate that progression through division (and the shutdown of division gene expression) depends on successful assembly of the ESCRT-III-based division ring, rather than ring constriction or membrane abscission. While the molecular basis by which cells monitor division ring assembly remains to be determined, this suggests the presence of a checkpoint-like feedback mechanism that holds cells in an active transcriptional state with TFB2 targets highly expressed until the division ring is complete. Since the assembly of the division ring is also required for chromosome segregation (*24*), ring assembly appears to be the major decision point in progression through D-phase. Since this occurs prior to the commitment to DNA segregation, membrane remodelling and the physical separation of daughter cells, the arrests occurs at a similar stage in the process of cell division as the spindle assembly checkpoint in eukaryotes, which monitors completion of the spindle as a prerequisite for triggering genome separation and cytokinesis.

These data lead us to propose a model for the regulation of cell division in *Sulfolobus*. As G2 cells enter D-phase, TFB2 is initially expressed at low levels and is subsequently amplified by autoactivation of its own promoter. Elevated TFB2 level then induces the expression of D-phase genes, including the *cdv* genes, along with the transcriptional repressor aCcr1 (which provides a delayed negative feedback). As Cdv proteins accumulate, a ring is assembled. Until this is complete, the ring assembly checkpoint maintains TFB2 expression. Following ring formation, TFB2 is rapidly degraded. Concomitantly, aCcr1, shuts off the division genes, including *tfb2* itself, allowing the transition into the next round of DNA replication. Importantly, the TFB2 regulon covers ∼30% of the previously identified cyclically expressed genes (*47*) (Supplementary data S2), suggesting that TFB2 is the main driver of the D-phase transcriptional program. The specific accumulation and repression dynamics of these genes can then be fine-tuned by repressors such as aCcrs and CCTF1 (*10*, *11*). In this sense, TFB2 plays a key role in driving the progression through D-phase, along with exit from division to allow subsequent entry into G1/S-phase.

As made clear in our study, despite involving many distinct molecular components, the archaeal and eukaryotic cell cycles share several regulatory principles including cyclic expression and degradation of key cell cycle regulators, coupling of positive and negative feedback circuits to generate transcriptional waves, and coordination of molecular and cell biological events through checkpoint systems. Our findings demonstrate that these regulatory principles can be implemented in an archaeal system without canonical CDK–cyclin machinery. The emergence of TFB2 as a dedicated cell-cycle regulator therefore illustrates how an ordered cell cycle can evolve through the repurposing and specialization of pre-existing transcriptional machinery and coupling to a cell cycle checkpoint, providing an alternative molecular solution to the problem of coordinating cell-cycle progression with gene expression.

## Acknowledgement

We would like to thank Dr. Arthur Radoux, Dr. Alice Cezanne and Dr. Fraser MacLeod for the fruitful discussion. We are grateful to Declan Barker for help with the aCcr1 purification. All flow cytometry and cell sorting experiments were performed at the Medical Research Council-Laboratory of Molecular Biology Flow Cytometry core facility, and we would like to thank members of the Flow Cytometry Facility for their technical support. The RNA-Seq library preps and sequencings were performed by CRUK Cambridge Institute Genomics Core, and we would like to thank members of the Genomics and Bioinformatics Core for their technical support. The RNA-Seq and phylogenetic analyses were performed in part via the Galaxy.org and Galaxy.eu servers. Y.-W.K. was supported by an EMBO postdoctoral fellowship (ALTF 903-2021) and by the Medical Research Council-Laboratory of Molecular Biology (MC_UP_1201/27); F.S. received support from the German Research Foundation (DFG Graduate Training Program 1431). F.B. and F.W. are supported by the Wellcome Trust (WT310064/Z/24/Z) to F.W. B.B. received support for work in Sulfolobus from the Medical Research Council-Laboratory of Molecular Biology (MC_UP_1201/27), the Wellcome Trust (222460/Z/21/Z), and the Life Sciences Moore-Simons Foundation (735929LPI).

## Code availability

All code for the analysis of ChIP-seq data is available at github.com/fblombach/TFB2_ChIP-seq.

## Competing interests

The authors declare that they have no conflict of interest.

## Author contributions

Y.-W.K. and B.B. conceived the project; Y.-W.K. performed all experiments and phylogenetic analysis; F.B. performed all ChIP experiments and corresponding bioinformatics analyses; F.S. and B.S. provided the endogenously tagged TFB1 and TFB2 strains; B.B. and F.W. supervised the project; Y.-W.K., F.B., F.W and B.B. wrote the manuscript with input from all authors.

## Notes

### Competing Interest Statement

The authors have declared no competing interest.

## References

1. M. Fischer, A. E. Schade, T. B. Branigan, G. A. Müller, J. A. DeCaprio, Coordinating gene expression during the cell cycle. Trends Biochem. Sci. 47, 1009–1022 (2022).

2. C. Bertoli, J. M. Skotheim, R. A. M. de Bruin, Control of cell cycle transcription during G1 and S phases. Nat. Rev. Mol. Cell Biol. 14, 518–528 (2013).

3. S. J. Rahi, K. Pecani, A. Ondracka, C. Oikonomou, F. R. Cross, The CDK-APC/C Oscillator Predominantly Entrains Periodic Cell-Cycle Transcription. Cell 165, 475–487 (2016).

4. A. Cezanne, S. Foo, Y.-W. Kuo, B. Baum, The Archaeal Cell Cycle. Annu. Rev. Cell Dev. Biol. 40, 1–23 (2024).

5. A.-C. Lindås, R. Bernander, The cell cycle of archaea. Nat. Rev. Microbiol. 11, 627–638 (2013).

6. M. Lundgren, R. Bernander, Genome-wide transcription map of an archaeal cell cycle. Proc. Natl. Acad. Sci. U. S. A. 104, 2939–2944 (2007).

7. M. V. Gomez-Raya-Vilanova, M. Krupovic, Regulation of eukaryotic-like cell cycle progression in archaea is coming into focus. Proc. Natl. Acad. Sci. U. S. A. 122, e2528525122 (2025).

8. X. Li, L.-M. Cristina, M.-A. Laura, P. Xu, A clade of RHH proteins ubiquitous in Sulfolobales and their viruses regulates cell cycle progression. Nucleic Acids Res. 51, 1724–1739 (2023).

9. Y. Yang, J. Liu, X. Fu, F. Zhou, S. Zhang, X. Zhang, Q. Huang, M. Krupovic, Q. She, J. Ni, Y. Shen, A novel RHH family transcription factor aCcr1 and its viral homologs dictate cell cycle progression in archaea. Nucleic Acids Res. 51, 1707–1723 (2023).

10. Y. Yang, S. Liang, Z. Geng, M. V. Gomez-Raya-Vilanova, W. Xia, J. Liu, Q. Huang, J. Ni, Q. She, M. Krupovic, Y. Shen, Successive waves of transcriptional repression and de-repression license cell cycle progression in an archaeon. Nucleic Acids Res. 54, gkag526 (2026).

11. Y.-W. Kuo, J. Traparić, S. Foo, B. Baum, The mechanism of cell-cycle-dependent proteasomal degradation of archaeal ESCRT-III homolog CdvB in Sulfolobus. EMBO J., doi: 10.1038/s44318-025-00688-7 (2026).

12. N. Takemata, R. Y. Samson, S. D. Bell, Physical and Functional Compartmentalization of Archaeal Chromosomes. Cell 179, 165–179.e18 (2019).

13. M. B. Elowitz, S. Leibler, A synthetic oscillatory network of transcriptional regulators. Nature 403, 335–338 (2000).

14. J. E. Ferrell, T. Y.-C. Tsai, Q. Yang, Modeling the cell cycle: why do certain circuits oscillate? Cell 144, 874–885 (2011).

15. J. R. Pomerening, S. Y. Kim, J. E. Ferrell, Systems-level dissection of the cell-cycle oscillator: bypassing positive feedback produces damped oscillations. Cell 122, 565–578 (2005).

16. J. Stricker, S. Cookson, M. R. Bennett, W. H. Mather, L. S. Tsimring, J. Hasty, A fast, robust and tunable synthetic gene oscillator. Nature 456, 516–519 (2008).

17. A. R. Araujo, L. Gelens, R. S. M. Sheriff, S. D. M. Santos, Positive Feedback Keeps Duration of Mitosis Temporally Insulated from Upstream Cell-Cycle Events. Mol. Cell 64, 362–375 (2016).

18. E. He, O. Kapuy, R. A. Oliveira, F. Uhlmann, J. J. Tyson, B. Novák, System-level feedbacks make the anaphase switch irreversible. Proc. Natl. Acad. Sci. U. S. A. 108, 10016–10021 (2011).

19. S. López-Avilés, O. Kapuy, B. Novák, F. Uhlmann, Irreversibility of mitotic exit is the consequence of systems-level feedback. Nature 459, 592–595 (2009).

20. G. Charvin, C. Oikonomou, E. D. Siggia, F. R. Cross, Origin of irreversibility of cell cycle start in budding yeast. PLoS Biol. 8, e1000284 (2010).

21. R. Bernander, A. Poplawski, Cell cycle characteristics of thermophilic archaea. J. Bacteriol. 179, 4963–4969 (1997).

22. G. Tarrason Risa, F. Hurtig, S. Bray, A. E. Hafner, L. Harker-Kirschneck, P. Faull, C. Davis, D. Papatziamou, D. R. Mutavchiev, C. Fan, L. Meneguello, A. Arashiro Pulschen, G. Dey, S. Culley, M. Kilkenny, D. P. Souza, L. Pellegrini, R. A. M. de Bruin, R. Henriques, A. P. Snijders, A. Šarić, A.-C. Lindås, N. P. Robinson, B. Baum, The proteasome controls ESCRT-III–mediated cell division in an archaeon. Science 369, eaaz2532 (2020).

23. A. Cezanne, B. Hoogenberg, B. Baum, Probing archaeal cell biology: exploring the use of dyes in the imaging of Sulfolobus cells. Front. Microbiol. 14, 1233032 (2023).

24. J. Parham, V. Sorichetti, A. Cezanne, S. Foo, Y.-W. Kuo, B. Hoogenberg, A. Radoux-Mergault, E. Mawdesley, L. D. Gatward, J. Boulanger, U. Schulze, A. Šarić, B. Baum, Temporal and spatial coordination of DNA segregation and cell division in an archaeon. Proc. Natl. Acad. Sci. U. S. A. 122, e2513939122 (2025).

25. J. McCullough, A. Frost, W. I. Sundquist, Structures, Functions, and Dynamics of ESCRT-III/Vps4 Membrane Remodeling and Fission Complexes. Annu. Rev. Cell Dev. Biol. 34, 85–109 (2018).

26. F. Hurtig, T. C. Q. Burgers, A. Cezanne, X. Jiang, F. N. Mol, J. Traparić, A. A. Pulschen, T. Nierhaus, G. Tarrason-Risa, L. Harker-Kirschneck, J. Löwe, A. Šarić, R. Vlijm, B. Baum, The patterned assembly and stepwise Vps4-mediated disassembly of composite ESCRT-III polymers drives archaeal cell division. Sci. Adv. 9, eade5224 (2023).

27. J. Liu, M. Lelek, Y. Yang, A. Salles, C. Zimmer, Y. Shen, M. Krupovic, A relay race of ESCRT-III paralogs drives cell division in a hyperthermophilic archaeon. mBio 16, e0099124 (2025).

28. R. Y. Samson, T. Obita, B. Hodgson, M. K. Shaw, P. L.-G. Chong, R. L. Williams, S. D. Bell, Molecular and structural basis of ESCRT-III recruitment to membranes during archaeal cell division. Mol. Cell 41, 186–196 (2011).

29. M. V. Gomez-Raya-Vilanova, J. Teulière, S. Medvedeva, Y. Dai, E. Corel, P. Lopez, F.-J. Lapointe, D. Bhattacharya, L.-P. Haraoui, E. Turc, M. Monot, V. Cvirkaite-Krupovic, E. Bapteste, M. Krupovic, Transcriptional landscape of the cell cycle in a model thermoacidophilic archaeon reveals similarities to eukaryotes. Nat. Commun. 16, 5697 (2025).

30. K. S. Makarova, Y. I. Wolf, E. V. Koonin, Archaeal Clusters of Orthologous Genes (arCOGs): An Update and Application for Analysis of Shared Features between Thermococcales, Methanococcales, and Methanobacteriales. Life 5, 818–840 (2015).

31. L. Hoffmann, A. Schummer, J. Reimann, M. F. Haurat, A. J. Wilson, M. Beeby, B. Warscheid, S.-V. Albers, Expanding the archaellum regulatory network - the eukaryotic protein kinases ArnC and ArnD influence motility of Sulfolobus acidocaldarius. MicrobiologyOpen 6, e00414 (2017).

32. Y. Wu, Q. Gan, K. Ning, R. Zhang, P. Wu, X. Feng, Q. She, J. Ni, Y. Shen, Q. Huang, Phosphorylation of the α subunit inhibits proteasome assembly and regulates cell cycle in an archaeon. Curr. Biol. CB 36, 979–994.e6 (2026).

33. Y. Xu, T. Gristwood, B. Hodgson, J. C. Trinidad, S.-V. Albers, S. D. Bell, Archaeal orthologs of Cdc45 and GINS form a stable complex that stimulates the helicase activity of MCM. Proc. Natl. Acad. Sci. U. S. A. 113, 13390–13395 (2016).

34. S. Lang, L. Huang, The Sulfolobus solfataricus GINS Complex Stimulates DNA Binding and Processive DNA Unwinding by Minichromosome Maintenance Helicase. J. Bacteriol. 197, 3409–3420 (2015).

35. Y. Yang, S. Liang, J. Liu, X. Fu, P. Wu, H. Li, J. Ni, Q. She, M. Krupovic, Y. Shen, Cran1, member of a new class of OLD family ATPases, functions in cell cycle progression in an archaeon. EMBO Rep. 27, 208–229 (2026).

36. L. Lemmens, K. Wang, E. Ruykens, V. T. Nguyen, A.-C. Lindås, R. Willaert, M. Couturier, E. Peeters, DNA-Binding Properties of a Novel Crenarchaeal Chromatin-Organizing Protein in Sulfolobus acidocaldarius. Biomolecules 12, 524 (2022).

37. I. G. Duggin, N. Dubarry, S. D. Bell, Replication termination and chromosome dimer resolution in the archaeon Sulfolobus solfataricus. EMBO J. 30, 145–153 (2011).

38. F. Werner, D. Grohmann, Evolution of multisubunit RNA polymerases in the three domains of life. Nat. Rev. Microbiol. 9, 85–98 (2011).

39. J. D. Parvin, P. A. Sharp, DNA topology and a minimal set of basal factors for transcription by RNA polymerase II. Cell 73, 533–540 (1993).

40. J. A. Coker, S. DasSarma, Genetic and transcriptomic analysis of transcription factor genes in the model halophilic Archaeon: coordinate action of TbpD and TfbA. BMC Genet. 8, 61 (2007).

41. C. Petitjean, K. S. Makarova, Y. I. Wolf, E. V. Koonin, Extreme Deviations from Expected Evolutionary Rates in Archaeal Protein Families. Genome Biol. Evol. 9, 2791–2811 (2017).

42. E. A. Pelve, A.-C. Lindås, W. Martens-Habbena, J. R. de la Torre, D. A. Stahl, R. Bernander, Cdv-based cell division and cell cycle organization in the thaumarchaeon Nitrosopumilus maritimus. Mol. Microbiol. 82, 555–566 (2011).

43. F. Blombach, T. Fouqueau, D. Matelska, K. Smollett, F. Werner, Promoter-proximal elongation regulates transcription in archaea. Nat. Commun. 12, 5524 (2021).

44. C. Zhang, A. P. R. Phillips, R. L. Wipfler, G. J. Olsen, R. J. Whitaker, The essential genome of the crenarchaeal model Sulfolobus islandicus. Nat. Commun. 9, 4908 (2018).

45. S. Dexl, R. Reichelt, K. Kraatz, S. Schulz, D. Grohmann, M. Bartlett, M. Thomm, Displacement of the transcription factor B reader domain during transcription initiation. Nucleic Acids Res. 46, 10066–10081 (2018).

46. R. Dhanaraju, R. Y. Samson, X. Feng, A. Costa, G. Gonzalez-Gutierrez, S. D. Bell, An archaeal nucleoid-associated protein binds an essential motif in DNA replication origins. Nat. Commun. 16, 5230 (2025).

47. M. Lundgren, L. Malandrin, S. Eriksson, H. Huber, R. Bernander, Cell cycle characteristics of crenarchaeota: unity among diversity. J. Bacteriol. 190, 5362–5367 (2008).

48. T. L. Bailey, C. Elkan, Fitting a mixture model by expectation maximization to discover motifs in biopolymers. Proc. Int. Conf. Intell. Syst. Mol. Biol. 2, 28–36 (1994).

49. S. Turkarslan, D. J. Reiss, G. Gibbins, W. L. Su, M. Pan, J. C. Bare, C. L. Plaisier, N. S. Baliga, Niche adaptation by expansion and reprogramming of general transcription factors. Mol. Syst. Biol. 7, 554 (2011).

50. F. Blombach, D. Grohmann, Same same but different: The evolution of TBP in archaea and their eukaryotic offspring. Transcription 8, 162–168 (2017).

51. A. Klotzbücher, E. Stewart, D. Harrison, T. Hunt, The “destruction box” of cyclin A allows B-type cyclins to be ubiquitinated, but not efficiently destroyed. EMBO J. 15, 3053–3064 (1996).

52. V. Ramachandran, M. Matzkies, A. Dienemann, F. Sprenger, Cyclin A degradation employs preferentially used lysines and a cyclin box function other than Cdk1 binding. Cell Cycle 6, 171–181 (2007).

53. M. E. Noble, J. A. Endicott, N. R. Brown, L. N. Johnson, The cyclin box fold: protein recognition in cell-cycle and transcription control. Trends Biochem. Sci. 22, 482–487 (1997).

54. T. J. Gibson, J. D. Thompson, A. Blocker, T. Kouzarides, Evidence for a protein domain superfamily shared by the cyclins, TFIIB and RB/p107. Nucleic Acids Res. 22, 946–952 (1994).

55. L. Aravind, V. Anantharaman, S. Balaji, M. M. Babu, L. M. Iyer, The many faces of the helix-turn-helix domain: transcription regulation and beyond. FEMS Microbiol. Rev. 29, 231–262 (2005).

56. J. Kato, H. Matsushime, S. W. Hiebert, M. E. Ewen, C. J. Sherr, Direct binding of cyclin D to the retinoblastoma gene product (pRb) and pRb phosphorylation by the cyclin D-dependent kinase CDK4. Genes Dev. 7, 331–342 (1993).

57. B. L. Allen, D. J. Taatjes, The Mediator complex: a central integrator of transcription. Nat. Rev. Mol. Cell Biol. 16, 155–166 (2015).

58. R. Shiekhattar, F. Mermelstein, R. P. Fisher, R. Drapkin, B. Dynlacht, H. C. Wessling, D. O. Morgan, D. Reinberg, Cdk-activating kinase complex is a component of human transcription factor TFIIH. Nature 374, 283–287 (1995).

59. S. Foo, Y.-W. Kuo, J. Traparić, D. W. Grogan, B. Baum, A temperature-sensitive mutant screen reveals a translational stress-induced cell-cycle arrest in a thermophilic archaeon. Mol. Biol. Cell 36, ar106 (2025).

60. M. Wagner, M. van Wolferen, A. Wagner, K. Lassak, B. H. Meyer, J. Reimann, S.-V. Albers, Versatile Genetic Tool Box for the Crenarchaeote Sulfolobus acidocaldarius. Front. Microbiol. 3, 214 (2012).

61. N. van der Kolk, A. Wagner, M. Wagner, B. Waßmer, B. Siebers, S.-V. Albers, Identification of XylR, the Activator of Arabinose/Xylose Inducible Regulon in Sulfolobus acidocaldarius and Its Application for Homologous Protein Expression. Front. Microbiol. 11, 1066 (2020).

62. X. Ye, A. Recalde, S.-V. Albers, M. van Wolferen, Methods for Markerless Gene Deletion and Plasmid-Based Expression in Sulfolobus acidocaldarius. Methods Mol. Biol. 2522, 135–144 (2022).

63. J. Schindelin, I. Arganda-Carreras, E. Frise, V. Kaynig, M. Longair, T. Pietzsch, S. Preibisch, C. Rueden, S. Saalfeld, B. Schmid, J.-Y. Tinevez, D. J. White, V. Hartenstein, K. Eliceiri, P. Tomancak, A. Cardona, Fiji: an open-source platform for biological-image analysis. Nat. Methods 9, 676–682 (2012).

64. Galaxy Community, Galaxy for accessible, reproducible, and collaborative data analyses: 2026 update. Nucleic Acids Res., gkag469 (2026).

65. A. Kechin, U. Boyarskikh, A. Kel, M. Filipenko, cutPrimers: A New Tool for Accurate Cutting of Primers from Reads of Targeted Next Generation Sequencing. J. Comput. Biol. J. Comput. Mol. Cell Biol. 24, 1138–1143 (2017).

66. H. Li, R. Durbin, Fast and accurate short read alignment with Burrows-Wheeler transform. Bioinformatics 25, 1754–1760 (2009).

67. B. Langmead, S. L. Salzberg, Fast gapped-read alignment with Bowtie 2. Nat. Methods 9, 357–359 (2012).

68. Y. Liao, G. K. Smyth, W. Shi, featureCounts: an efficient general purpose program for assigning sequence reads to genomic features. Bioinformatics 30, 923–930 (2014).

69. M. I. Love, W. Huber, S. Anders, Moderated estimation of fold change and dispersion for RNA-seq data with DESeq2. Genome Biol. 15, 550 (2014).

70. F. Blombach, K. L. Smollett, F. Werner, ChIP-Seq Occupancy Mapping of the Archaeal Transcription Machinery. Methods Mol. Biol. 2522, 209–222 (2022).

71. O. Cohen, S. Doron, O. Wurtzel, D. Dar, S. Edelheit, I. Karunker, E. Mick, R. Sorek, Comparative transcriptomics across the prokaryotic tree of life. Nucleic Acids Res. 44, W46–53 (2016).

72. Q. Li, J. B. Brown, H. Huang, P. J. Bickel, Measuring reproducibility of high-throughput experiments. Ann. Appl. Stat. 5 (2011).

73. S. R. A. Fisher, CONFIDENCE LIMITS FOR A CROSS-PRODUCT RATIO. Aust. J. Stat. 4, 41–41 (1962).

74. R. D. Finn, J. Clements, S. R. Eddy, HMMER web server: interactive sequence similarity searching. Nucleic Acids Res. 39, W29–37 (2011).

75. L. Fu, B. Niu, Z. Zhu, S. Wu, W. Li, CD-HIT: accelerated for clustering the next-generation sequencing data. Bioinformatics 28, 3150–3152 (2012).

76. S. Capella-Gutiérrez, J. M. Silla-Martínez, T. Gabaldón, trimAl: a tool for automated alignment trimming in large-scale phylogenetic analyses. Bioinformatics 25, 1972–1973 (2009).

77. L.-T. Nguyen, H. A. Schmidt, A. von Haeseler, B. Q. Minh, IQ-TREE: a fast and effective stochastic algorithm for estimating maximum-likelihood phylogenies. Mol. Biol. Evol. 32, 268–274 (2015).

78. D. T. Hoang, O. Chernomor, A. von Haeseler, B. Q. Minh, L. S. Vinh, UFBoot2: Improving the Ultrafast Bootstrap Approximation. Mol. Biol. Evol. 35, 518–522 (2018).

79. S. Kalyaanamoorthy, B. Q. Minh, T. K. F. Wong, A. von Haeseler, L. S. Jermiin, ModelFinder: fast model selection for accurate phylogenetic estimates. Nat. Methods 14, 587–589 (2017).

80. I. Letunic, P. Bork, Interactive Tree of Life (iTOL) v6: recent updates to the phylogenetic tree display and annotation tool. Nucleic Acids Res. 52, W78–W82 (2024).

