## Supplemental materials for "A TFIIB paralog drives cyclic transcription to orchestrate the archaeal cell cycle"

### Materials and Methods

#### Cell culture and induction

All *Sulfolobus acidocaldarius* strains were cultured in Brock medium supplemented with 0.1% NZ-amine, 0.2% sucrose, 0.04 g/L $FeCl_3(H_2O)_6$ (denoted as BNS media here on) and the pH was adjusted to 3 with 50% (v/v) sulfuric acid as previously described (*59*). The cultures were grown at 75 °C with 160 rpm shaking. 4 mg/L of uracil was supplemented for culturing of uracil auxotrophic strain MW001 (*60*), and endogenously tagged TFB1 and TFB2 strains. All cultures were passed at least once after inoculation from the frozen glycerol stock for all experiments. Induction of the arabinose-inducible promoter was performed by addition of 0.2% L-arabinose at the early log-phase ($OD_{600}$~0.1-0.2).

#### Molecular genetics and plasmid transformation

To construct Vps4 and aCcr1 overexpression plasmids, the pSVAaraFX plasmid vector containing the arabinose-inducible promoter (*61*) was double digested by the restriction enzymes NcoI and XhoI, followed by gel extraction. The coding sequence of *S. acidocaldarius* Vps4 gene (*saci_1372*) and aCcr1 gene (*saci_0942*) were amplified from the genomic DNA of wildtype cells by PCR using primers containing flanking sequences that overlap with the digested vector and assembled using Gibson assembly. Arabinose-inducible TFB2-FLAG-StrepII plasmid was constructed similarly with the coding sequence amplified from the genomic DNA of the TFB2 endogenously tagged strain. Truncation constructs of TFB2 were then generated by using Q5-site directed mutagenesis kit. The aCcr1/TFB2-FLAG-StrepII co-expression plasmid was generated by first linearising the pSVAaraFX-TFB2-FLAG-StrepII plasmid by PCR and the upstream sequence (16 nucleotides) of the endogenous TFB2 gene was added by flanking the forward primer. The aCcr1 coding sequence was then PCR amplified from the wildtype genomic DNA using primers containing flanking sequences that overlap with the two ends of the linearized vector and assembled by Gibson assembly. All cloned plasmids were verified by Sanger sequencing.

Cloned plasmids were transformed into *E. coli* ER1821 strain that contains pM.EsaBC41 plasmid for cytosine methylation. The methylated plasmids were then extracted and transformed into uracil auxotrophic strain (MW001 unless otherwise noted) by electroporation. The electroporated cells were immediately resuspended in 1 mL of recovery medium (Brock media without supplementation of additional NZ-amine and $FeCl_3$ solution, pH ~5), and incubated at 75 °C for 0.5-1 hr with shaking. The recovered cells were then plated on Brock medium plates containing 0.6% gelrite (*62*) without uracil, and incubated at 75 °C for 5 to 7 days. Colonies were then picked and grown in BNS media at 75 °C to stationary phase, and verified by PCR genotyping and Sanger sequencing of the PCR product. Positive clone cultures were then centrifuged at 4300xg for 4 min. The cell pellet was resuspended in Brock media with 50% glycerol (without additional sulfuric acid, pH ~5) and stored as a frozen glycerol stock at -70 °C.

The markerless endogenously tagged TFB1 (saci_0866) and TFB2 (saci_1341) were generated using the homologous recombination-based pop in/pop out strategy in the MW001 background as previously described (*60*) to introduce a C-terminal FLAG-StrepII tag with a Gly-Ala linker at the endogenous locus. Successfully tagged clones were identified by PCR genotyping followed by Sanger sequencing targeting the two loci.

### Synchronisation of *S. acidocaldarius*

*S. acidocaldarius* cultures were first arrested at G2 phase by addition of 2 mM acetic acid for 4.5 hr at 75 °C. The arrested cells were then washed by fresh media two times and resuspended in fresh media with ~80% of original culture volume to account for the cell loss during the washing steps. The released cultures were grown at 75 °C with 160 rpm shaking. For synchronisation of HA-TFB2ΔN strains, 0.2% arabinose was added into the cultures (including the empty vector controls) 2 hours before release, and 0.2% arabinose was supplemented in the washes and the release culture media. For synchronisation of uracil auxotrophic strains, 4 mg/L uracil was supplemented in all steps.

### Immunostaining and flow cytometry

To fix the cells for immunostaining, cold ethanol was added to the cell culture stepwise to a final concentration of 33%, 50%, and 70% with 5 min intervals on ice, followed by storage at 4 °C. To perform immunostaining, ~1.5 mL of fixed cells was centrifuged at 8000 x g, 3 min at room temperature to remove the ethanol, followed by two washes of phosphate buffer saline supplemented with 0.2% Tween-20 and 3% bovine serum albumin (hereafter denoted as PBSTA). The washed cells were then incubated with primary antibodies in PBSTA overnight at room temperature with 500 rpm shaking. The cells were then washed three times with PBST (without bovine serum albumin), and incubated with fluorophore-conjugated secondary antibodies in PBSTA for 1.5-2.5 hr at room temperature with shaking. For immunofluorescence imaging, 5 µg/mL AlexaFluor 647-conjugated concanavalin A (Invitrogen) was added along with the secondary antibodies. The stained cells were then washed with PBST for three times and resuspended in PBST supplemented with 2 µM Hoechst 33342 (Thermo Scientific) for flow cytometric analysis or 2 µM DAPI (Invitrogen) for immunofluorescence microscopy to stain the DNA.

All flow cytometry analyses were performed on a BD Biosciences LSRFortessa with excitation laser lines of 355, 488, 561 and 640 nm in conjunction with the emission filters 450/50, 530/30, 585/15 and 670/15, respectively. The Hoechst channel (355 nm excitation with emission filter 450/50 nm) was used for thresholding to identify immunostained cells and $2.5 \times 10^5$ events were recorded for each experiment unless otherwise noted. All flow cytometry data were analysed by FlowJo v10 software, and single cells were selected by gating the diagonal population in the area vs height plot in the Hoechst channel. Quantification of fluorescence intensity and proportions of cells in indicated gates was performed only on the selected single cells using FlowJo.

### Immunofluorescence microscopy

For immunofluorescence imaging, the stained cells were centrifuged onto a LabTek chamber (ThermoFisher) coated with 1% polyethyleneimine or 1% poly-L-lysine solution (750 x g, 1 hr at room temperature) and imaged on a Nikon Eclipse Ti2 inverted microscope equipped with a Yokogawa SoRa scanner unit and Prime 95B scientific complementary metal-oxide semiconductor (sCMOS) camera (Photometrics). Images were acquired by an oil immersion objective (Apo TIRF 100X/NA=1.49, Nikon) with an additional 2.8x magnification lens from the SoRa unit and ten z-sections were collected with a step of 0.22 µm. The exposure time of protein labels was set to 50 to 100 ms, with the DNA staining channel set to 500 ms and the laser intensities adjusted to prevent pixel saturation.

### Western blotting

Cells were lysed in the 1x Laemmli buffer (Biorad) and incubated at 98 °C for 10 min and centrifuged at 20,000 x g for 5 min at room temperature to remove insoluble cell debris. SDS-PAGE was then performed by loading the samples on a NuPAGE 4-12% Bis-Tris mini gel (Invitrogen) and running at 150 V with MES-SDS running buffer. The proteins were then transferred to a nitrocellulose membrane with 100 V for 1 hr at 4 °C. Membranes were blocked with 5% milk in PBS supplemented with 0.2% Tween-

20 (PBST) for 1 hr at room temperature, followed by incubation with primary antibodies in the blocking buffer overnight at 4 °C. The blots were then washed three times with PBST and incubated with the fluorophore-conjugated secondary antibodies in the dark for 2 hr at room temperature. The blots were washed with PBST for another three times and imaged using the Bio-Rad ChemiDoc system. The band intensities were quantified using the gel quantification function in Fiji (*63*).

RT-qPCR analysis of FACS sorted cells

To enrich the dividing cells, wildtype *S. acidocaldarius* (DSM639) was synchronised and ethanol fixed after release from acetic acid arrest and stored in -20 °C until the day before FACS sorting. The earliest timepoint with emerging cells with 1N DNA content was used for sorting (typically at 80-100 minutes after release from acetic acid arrest). This ensured that the cells with 1N DNA content are predominantly in G1 phase. The fixed cells were washed three times with cold PBSTA, and stained with anti-CdvB (1:400) and anti-CdvB2 (1:400) antibodies supplemented with 1U/μL RNase inhibitor Superase IN (Thermofisher) in PBSTA at 16 °C overnight with constant shaking. The cells were then washed three times with cold PBST and stained with fluorophore-conjugated secondary antibodies with 1U/μL Superase IN in PBSTA at 16 °C for 1.5 hr. The cells were further washed twice with cold PBST and resuspended in PBST containing 2 μM Hoechst 33342 DNA dye and 0.2U/μL Superase IN. The resuspended cells were passed through the 70 μm cell strainer and sorted on ThermoFisher BigFoot and BD FACSAria Fusion with a 70 μm nozzle at 60 psi. Populations of cells were gated by a combination of DNA dye staining and CdvB/CdvB2 signal as shown in Fig. 1. The sorted cells were collected in RNAlater solution (Invitrogen), and were centrifuged at 5000 x g for 40 minutes at 4 °C. The supernatant was then carefully removed, and the pellets were resuspended in 1 to 2 mL of Trizol solution (Invitrogen) and stored in -20 °C until RNA extraction.

RNA was first extracted by chloroform and centrifuged at 12,000 x g for 15 minutes at 4 °C. The aqueous phase was then taken out, and the RNA was then extracted using the RNA clean & concentrator kit (Zymo) along with an on-column DNaseI digestion. The eluted RNA was then used for cDNA synthesis using the iScript cDNA synthesis kit (Biorad) following manufacturer's protocol. The RT-qPCR analysis was then performed using iTaq SYBR-green kit (Biorad) with 16S rRNA as the housekeeping gene. The relative RNA transcript abundance was then calculated as $2^{Ct(housekeeping)-Ct(target)}$ followed by normalization to the constricting phase.

RNA extraction and RT-qPCR

To extract RNA for bulk RT-qPCR analysis and RNA-Seq, 10-20 mL of culture was centrifuged at 4300 x g for 4 min at room temperature, and the cell pellets were resuspended and lysed in 750 μL Trizol reagent (Invitrogen). The total cellular RNA was then purified by chloroform extraction, isopropanol precipitation and washed with 70% ethanol following the manufacturer's protocol. The RNA pellets were then resuspended in 100 µL 1x TURBO DNase buffer, followed by addition of TURBO DNase to the final concentration of 0.04U/µL (Invitrogen) and incubated at 37 °C for 20 min. The DNase-treated RNA samples were then purified by RNeasy kit (Qiagen) and eluted with nuclease-free water. One-step RT-qPCR reactions were prepared with the Luna One-Step RT-qPCR kit (New England Biolabs) with ~6-15 ng of total RNA used per reaction, followed by the analysis on an Applied Biosystems ViiA7 real-time PCR system. The transcript abundance was quantified by $2^{Ct(housekeeping)-Ct(target)}$ using SecY (saci_0574) as the housekeeping gene. The primer pairs used for all RT-qPCR experiments are summarised in Supplementary Table S2.

RNA-Seq experiment and analysis

For wild type Vps4 and Vps4$^{E209Q}$ overexpression strains, the cultures were induced with 0.2% arabinose for 5 hr at 75 °C before harvesting for RNA extraction following the aforementioned process. For the aCcr1 overexpression and the corresponding vector only control strain, 3 hr arabinose-induction was used. The RNA integrity (RIN) was examined by Bioanalyzer using the Prokaryote total RNA pico kit (Agilent). Only samples with RIN > 8 were used for RNA-Seq experiments, and four biological replicates were used for each condition. The ribosomal RNA was depleted from the total RNA samples by using the Pan-Archaea ribopool rRNA depletion kit (siTOOLs Biotech) following the manufacturer's instructions. The libraries were then prepared by using the Watchmaker RNA Library Prep Kit with Polaris depletion (Watchmaker Genomics) but started after the depletion step in the manufacturer's protocol. 7 PCR cycles were used for the library amplification step and IDT xGen Stubby adapter UDIs were used. The library was spiked in with 750 pM with a 1% PhiX spike-in and sequenced on NextSeq2000 using the P1 reagent kit (paired-end 50 bp, Illumina).

All data processing after demultiplexing was performed on the Galaxy server (*64*). The sequencing data first underwent quality trimming using Cutadapt (*65*), and mapped to the *S. acidocaldarius* DSM639 genome (NCBI ASM1228v1) using Burrows-Wheeler Aligner (BWA) or Bowtie2 (*66*, *67*). Read counts corresponding to protein coding genes were quantified using featureCounts (*68*). The differential gene expression analyses were then performed by using DESeq2 (*69*).

*S. acidocaldarius* aCcr1 purification and antiserum preparation

The gene coding for aCcr1, *saci_0942*, was PCR-amplified using primers 5'-ggcggcatATGAGAGTAGTTACATTTAAGGTGGA and 5'-gcgcgctcgagAAGTTTGACTTTTTCTACCCTGGC, and cloned into pET21a+ vector (Novagen) via NdeI and XhoI restriction sites resulting in a C-terminal His$_6$-tag fusion. The sequence was verified by Sanger sequencing. Rosetta 2(DE3) cells (Novagen) were transformed by the resulting plasmid and heterologous expression was carried out for 4 hrs at 37 °C. The protein purification protocol was based on (*9*) using Ni-affinity purification on a His-trap column (Cytiva) and size exclusion chromatography on a Superdex 75 16/60 column (Cytiva). Purified protein was sent to Davids Biotechnologie (Regensburg, Germany) for rabbit antiserum production. Antibodies were purified by Protein G affinity-chromatography for ChIP experiments and quantified based on $A_{280nm}$ and the standard IgG extinction coefficient on a NanoDrop spectrophotometer (ThermoFisher).

ChIP-seq sample preparation and analysis

The ChIP-seq protocol was based on (*43*) and a detailed protocol is available here (*70*). For all ChIP-seq experiments, ~200 mL of culture was grown in a shaking incubator at 75 °C. The culture was transferred onto a hot plate maintained at the growth temperature right before crosslinked with 0.4% formaldehyde at 75 °C for 1 min with constant mixing using a magnetic stirrer. 1M Tris (pH 8.0) solution was then added to a final concentration of 100 mM for quenching. The culture was then cooled on ice for 15-30 minutes and centrifuged at 5000 x *g* for 10 min at 4 °C. The pellet was then resuspended in 15 mL of cold PBS and transferred to a 50 mL tube. The fixed cells were then centrifuged at 5000 x *g* for 5 minutes at 4 °C. The pellet was then further washed by 10 mL cold PBS followed by centrifugation at 5000 x g for 5 minutes at 4 °C twice. The pellet was then flash frozen by liquid nitrogen and stored in -70 °C until downstream sample processing.

The cross-linked cell pellet was thawed on ice and resuspended in ChIP lysis buffer (50 mM HEPES–NaOH pH 7.5, 140 mM NaCl, 1 mM EDTA, 0.1% sodium deoxycholate, 1% Triton X-100 with 1x protease inhibitor cocktail (Roche)). DNA was sheared to ~200bp average fragment size using a Qsonica Q700 cup sonicator. Lysate corresponding to 20 µg (Vps4 overexpression or 10 µg genomic DNA (synchronised cells, TFB2 overexpression, aCcr1 overexpression, based on $A_{260nm}$ absorption of purified chromatin input) was incubated overnight at 4 °C with 2 µg THE™ NWSHPQFEK Tag monoclonal antibody (Genscript) or polyclonal purified anti-aCcr1 antibody (see above). Antibodies were then captured with 50 µl Protein G Dynabeads (ThermoFisher) to isolate immuno-precipitated DNA. 1 ng of immuno-precipitated DNA or purified chromatin input was used for Illumina sequencing library preparation using NEBNext Ultra II DNA library prep kit for Illumina (New England Biolabs) without size selection step and 11 rounds of PCR amplification. Libraries were sequenced on a NovaSeq X platform (paired-end, 150 cycles). Sequencing quality was assessed using FastQC v0.12.1. The first 100 nt of reads were aligned to the *S. acidocaldarius* DSM639 genome using Bowtie2 (*67*) with the "—sensitive" preset plus "settings -3 50 –fr -X 1000". Bam file output was imported into the R environment (v4.3) using the Rsamtools 2.22.0 and rtracklayer 1.66.0 packages and sampled for the fragment sizes to match a normal distribution with mean 200 bp and standard deviation of 40 bp (see accompanying code) before export as bam files. These bam files formed the basis for any differential binding analysis and peak calling. For visualisation, the ChIP bam files were normalised against the respective chromatin input of each replicate using deeptools 3.5.6 bamCompare with signal extraction scaling based on 10,000 bins of 200 bp (Ramírez *et al*, 2016). All experiments were carried out with three biological replicates.

The ChIP-seq heatmap was generated using deeptools 3.5.6 computeMatrix and plotHeatmap based on primary TSSs (*71*) within 50 bp of TFB2 peaks. Where one TFB2 peak overlapped with two primary TSSs, the TSS with higher RNA-Seq expression levels was chosen.

<u>Peak calling of ChIP-seq</u>

Fragment size-adjusted bam files were used for peak calling using macs2 callpeak function with settings "-f BAM --nomodel -q 0.05 --keep-dup auto --call-summits" (v2.2.9.1) for each biological replicate. Data were imported into the R environment (v4.3) and pairwise reproducibility was assessed using the Cran IDR package (*72*) setting a local IDR threshold of 0.05. The final peak calling data set (optimal IDR peaks) and average enrichment was based on the biological replicate pair yielding the largest set of reproducible peaks in line with Encode Project guidelines. For the synchronised cell ChIP-seq data, we added the matching peaks from the other biological replicate for average enrichment calculation removing any peaks from the data set where no match in the other biological replicate could be found.

<u>Association of TFB2, TFB1 and aCcr1 with primary transcription start sites (TSSs) and start codons</u>

For the association of ChIP-seq peaks with the promoters of differentially expressed genes, we used the start codon as a proxy for the TSS where the primary TSS has not been determined. This is justified as most Sulfolobales genes have short 5' UTRs (*43*). Operon maps and ChIP-seq peak summit positions were imported into the R environment as GenomicRanges objects and association was tested using the findOverlaps function from the GenomicRanges package v1.58 with a maximal distance of 80 bp. Differentially expressed operons were defined as operons where at least one cistron shows differential expression in the corresponding RNA-Seq data set. Fisher's exact test was used to test association and calculate 95% confidence intervals according to (*73*). To avoid overcounting of TFB2/aCcr1 peak-associated transcription units (TUs) where a single ChIP-seq peak was associated with two divergent closely spaced TU starts, we counted only unique ChIP-seq peaks. To similarly estimate the number of peaks that would be expected to cover all TU starts lacking ChIP-seq peaks, we calculated weight factors for each TU start reflecting the likelihood that a ChIP-seq peak at this TU start would not overlap with a second TU start. The weight factors ranged from 0.5 (overlap) to 1 (no overlap) and were derived from a binominal regression models with the distance to the next TU start as explanatory variable and the overlap likelihood as dependent variable in R.

MEME motif discovery in TFB2 and aCcr1 bound regions

For aCcr1, ChIP-seq peaks with enrichment > 5 were selected (n = 135) and DNA regions flanking 50 bp on each site of the peak summit were used for the motif analysis. MEME v5.5.9 (*48*) was run with settings "-dna -mod anr -minw 6 -maxw 50 -objfun classic -revcomp -markov_order 0" allowing for any number of motifs per sequence to be found on both strands. MEME detected 227 sites within 129 peaks.

Phylogenetic analysis of TFB homologs

The initial set of TFB homologs was searched by phmmer (*74*) with a cutoff of E-value ≤ e-21 using the protein sequence of *S. acidocaldarius* TFB2 (saci_1341) as the query. Asgard archaeal TFB sequences were identified by BLAST using *S. acidocaldarius* TFB1 (saci_0866) and TFB2 as queries, respectively. The sequences were combined and filtered using cd-hit (*75*) with a cutoff of 85% sequence identity to remove redundant sequences. Multiple sequence alignment using MAFFT was then performed and heuristically trimmed by using trimAl (*76*). The sequence alignment was then used as the alignment input for hmmsearch from HMMER (*74*) against the RP15 database for eukaryotes and Reference Proteome for archaea and viruses, which were later combined with the Asgard archaeal TFB sequences identified by BLAST in NCBI clustered nr database. Sequences were then manually selected to remove sequences with uncertain/low-confidence annotations and overrepresented taxonomic groups. This gave a total of ~440 sequences.

Multiple sequence alignment with MAFFT was then performed on this collection of sequences using BLOSUM 62 matrix, and the alignment was trimmed by trimAl to remove sites containing a high proportion of gaps. The phylogenetic tree was then constructed using IQ-Tree (*77*) with ultrafast bootstrapping (*78*), using an automatic model search where a full tree search was invoked for higher accuracy (*79*). The phylogenetic trees were then visualised using iTOL (*80*). Note that maximum likelihood (ML) tree and consensus tree showed similar topology and only the ML tree was shown in Supplementary Figure S4.

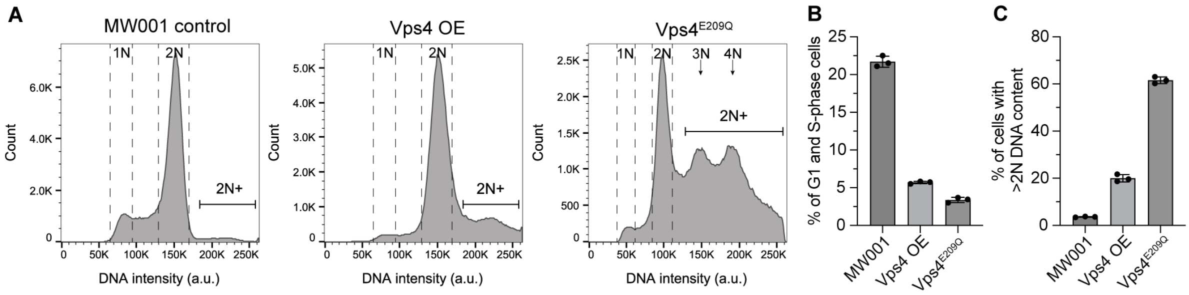


**Supplementary Figure S1. Overexpression of wildtype and dominant negative mutant of Vps4 disrupt Sulfolobus cell division. (A)** Example flow cytometry histograms of DNA content of asynchronized MW001 (background strain control), Vps4 overexpression (OE), and the dominant negative hydrolysis-deficient Vps4$^{E209Q}$ mutant (all with 5 hr arabinose induction). Note that the flow cytometer detector voltage was adjusted in the Vps4$^{E209Q}$ experiment to reduce extensive truncation of the population with high ploidy. **(B)** Percentage of cells in the G1 and S-phase in each condition quantified from flow cytometry analysis. **(C)** Percentage of cells with more than 2N DNA content (2N+) quantified from flow cytometry analysis. Error bars: mean±SD, N=3 biological replicates.

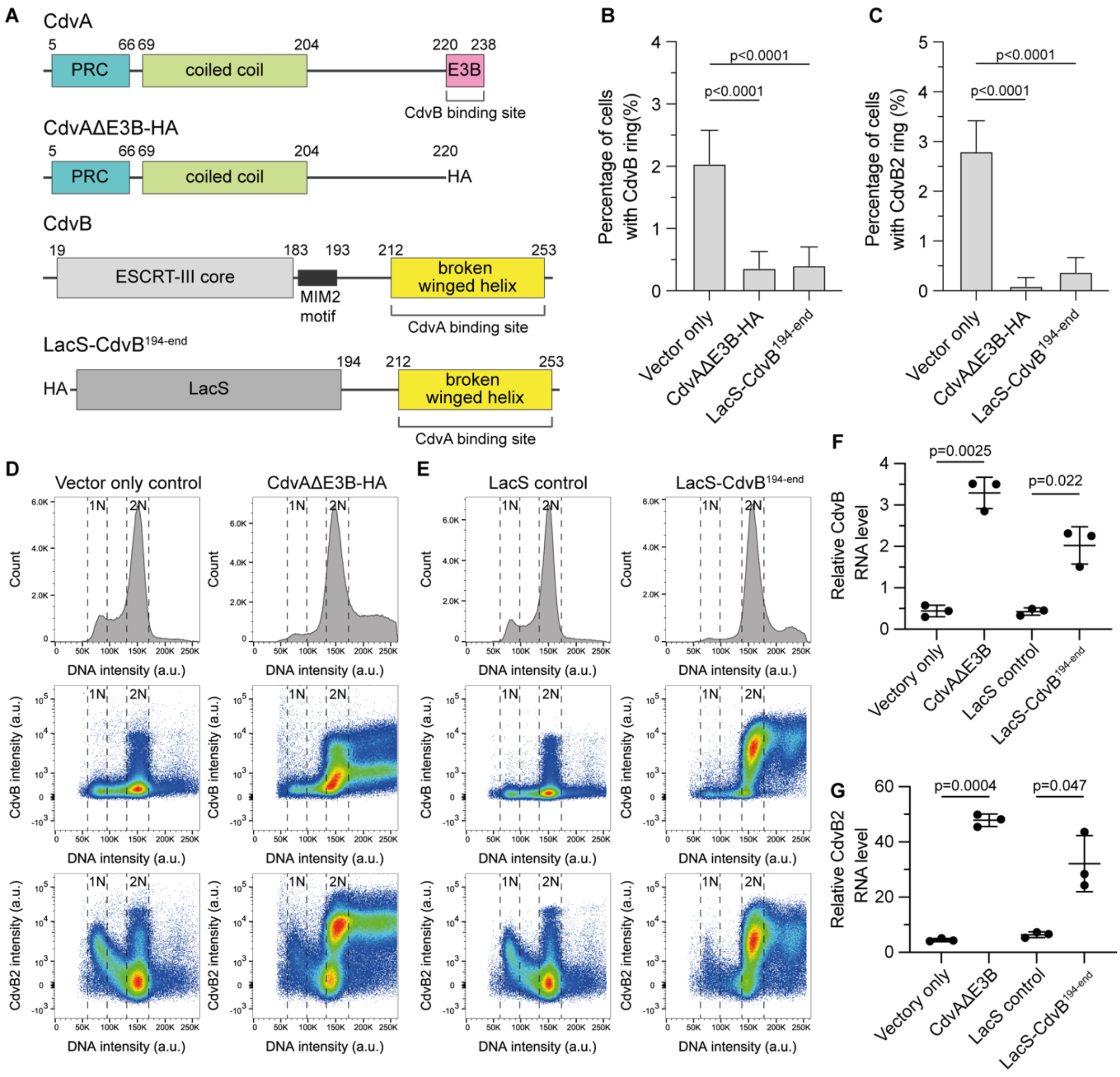


**Supplementary Figure S2. Perturbation of division ring assembly by disrupting interaction between CdvA and CdvB. (A)** Design of the CdvAΔE3B and LacS-CdvB$^{194\text{-end}}$ mutant. **(B)(C)** Percentage of CdvB and CdvB2 rings observed in vector only control, CdvAΔE3B and LacS-CdvB$^{194\text{-end}}$ expressing mutants from immunofluorescence imaging of ethanol-fixed cells. Error bars: 95% CI. Fisher's exact test, n=3288, 3312, 2965 cells pooled from 3 biological replicates each. **(D)(E)** Example flow cytometric histograms and scatter plots of CdvAΔE3B and LacS-CdvB$^{194\text{-end}}$ mutants with controls. Both mutants showed division defects indicated by a decrease of G1-S phase cells along with an increase in cells with >2N DNA content. Increase of CdvB and CdvB2 protein abundance was observed in the scatter plots. **(F)(G)** RT-qPCR analysis showed upregulation of CdvB and CdvB2 genes in both ring perturbation mutants (Welch t-test, error bars: mean±SD, N=3 biological replicates).

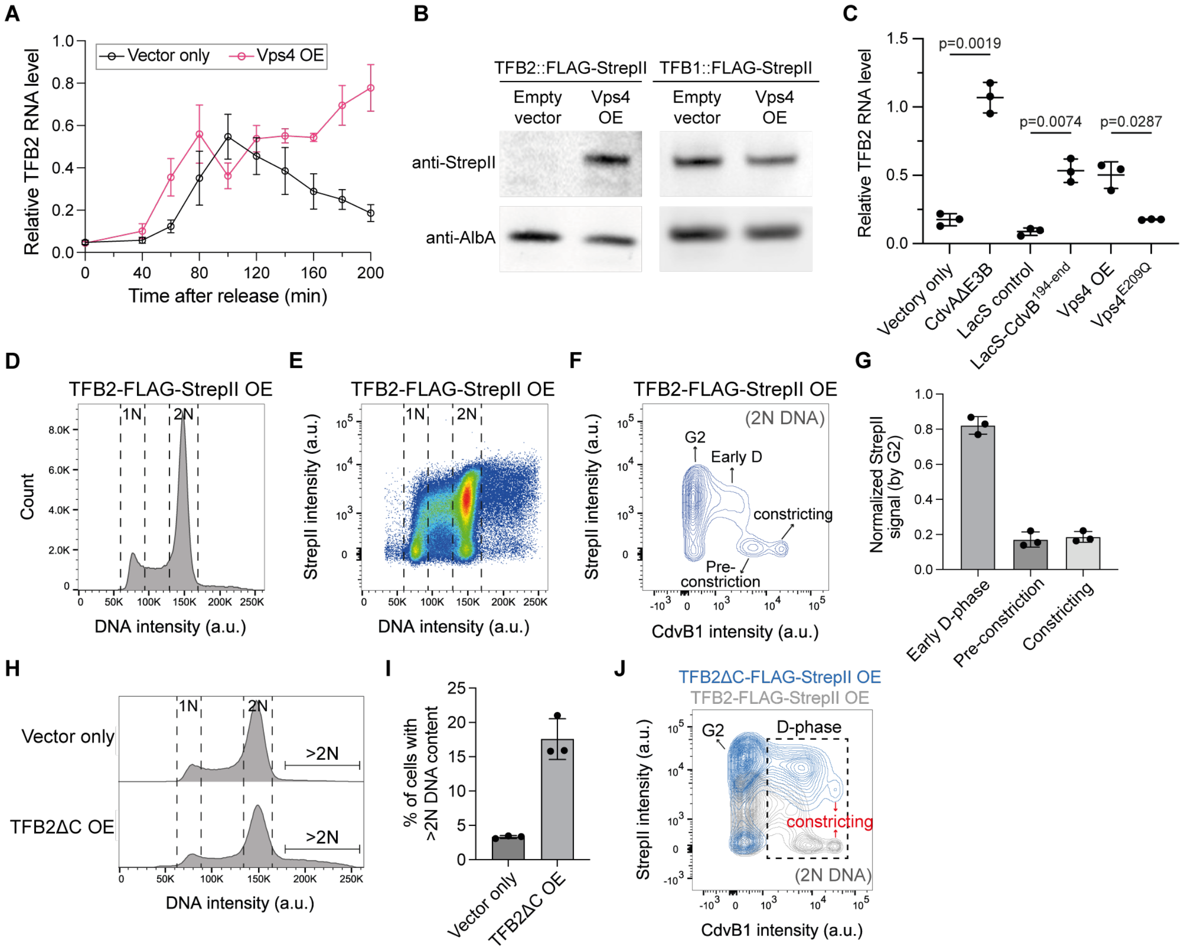


**Supplementary Figure S3. Expression level of TFB2 is cell cycle dependent. (A)** RT-qPCR analysis of synchronised vector only control and Vps4 OE cells released from acetic acid-induced arrest. TFB2 is cyclically expressed in vector only control, but continue to express in high level in the Vps4 OE cells. 0.2% arabinose were added 2 hr prior to release from acetic acid in both conditions. Error bars: mean±SEM. N=3 biological replicates. **(B)** Western blog of Vps4 overexpression in the endogenously tagged TFB2 (TFB2::FLAG-StrepII), and TFB1 (TFB1::FLAG-StrepII) strains showed that only TFB2 was upregulated when division ring assembly was perturbed. DNA binding protein AlbA was used as a loading control. **(C)** RT-qPCR analyses show that TFB2 is up-regulated in mutants that failed to assemble full division rings. Error bars: mean±SDs. Welch's t-test, N=3 biological replicates. **(D-F)** Flow cytometry histogram and scatter plot of TFB2-FLAG-StrepII overexpression cells (4 hr after arabinose induction). Cells with 2N DNA content was shown in (F). **(G)** Normalised average TFB2-FLAG-StrepII level (arabinose-induced) at different stage in D-phase quantified from flow cytometry analysis in (D-F). The substages were defined by DNA content, CdvB and CdvB1 signal intensity as described in Fig. 1. The StrepII signal was normalised by the average intensity in G2 phase. Error bars: mean±SD, N=3 biological replicates. **(H)** Flow cytometry histogram of cells expressing TFB2ΔC mutant (1-192aa) where the second cyclin box was truncated. **(I)** Quantification of the percentage of cells with >2N DNA content in (H). Error bars: mean±SD. **(J)** Flow cytometry scatter plot of TFB2ΔC expressing mutant with 2N DNA content (blue) overlaid with TFB2-FLAG-StrepII OE (grey) for reference. The substage assignment was based on CdvB1 and CdvB intensity as described in Fig. 1.

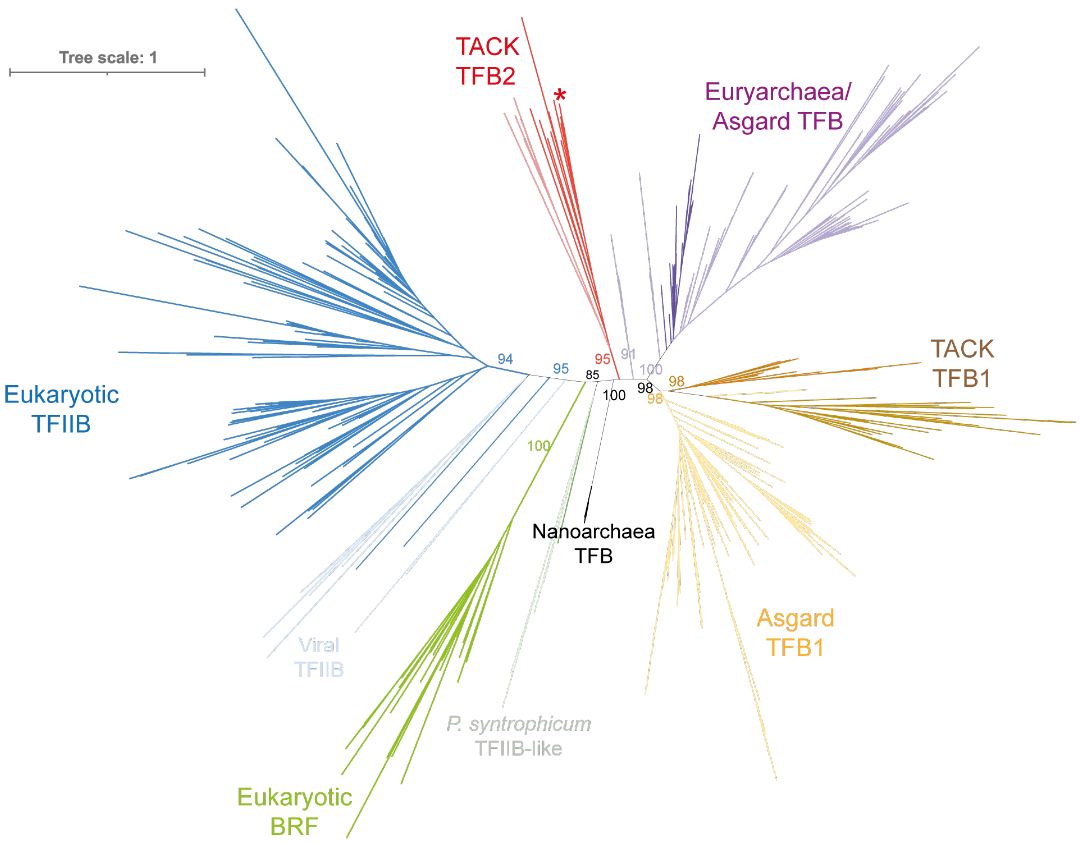


**Supplementary Fig. S4. Unrooted maximum likelihood tree of TFIIB homologues.** Different taxonomical groups were marked by various colours. *Sulfolobus acidocaldarius* TFB2 is labelled with asterisk. Support values are from IQ-TREE ultrafast bootstrapping.

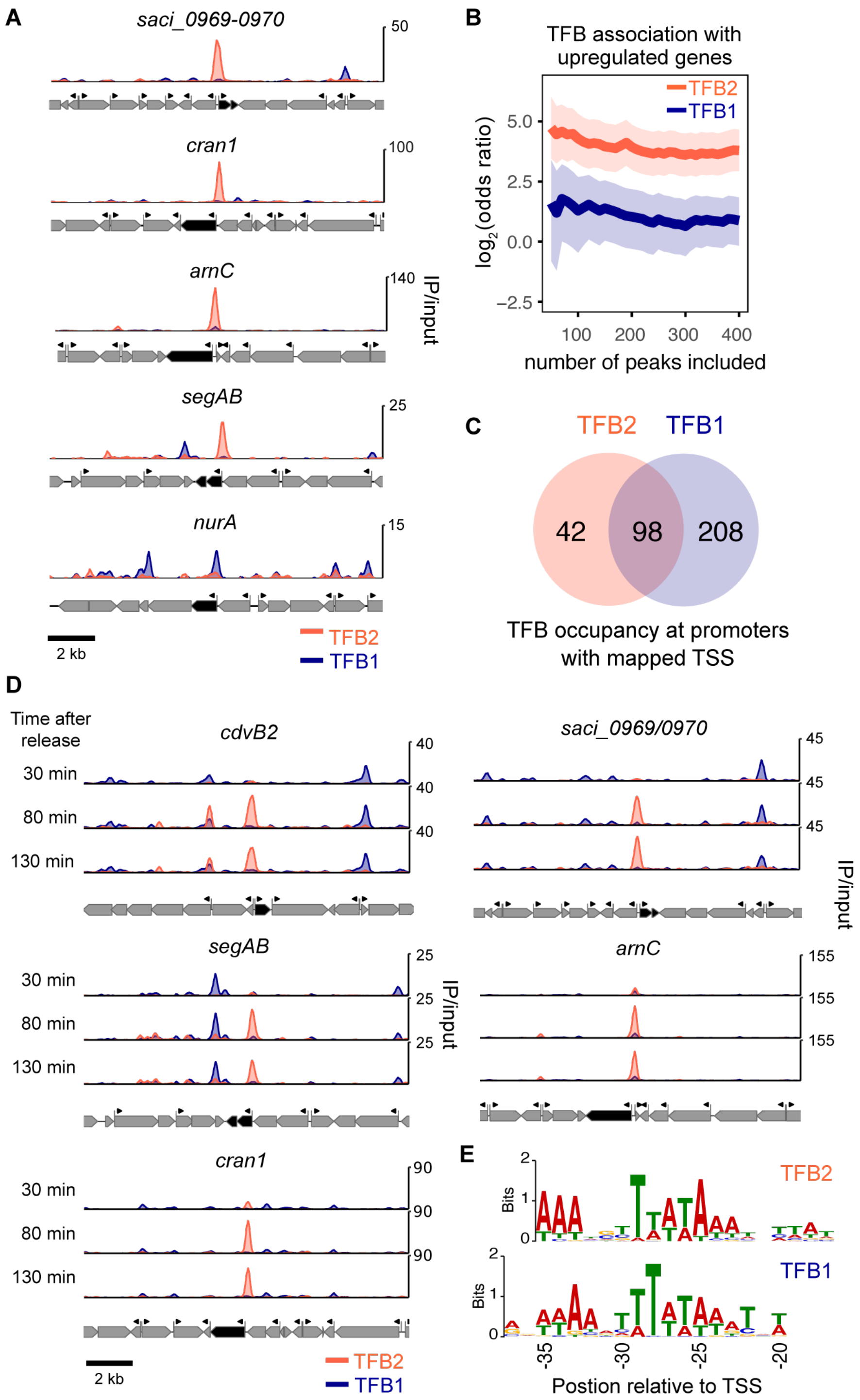


**Supplementary Figure S5. TFB2 binds to the promoter region of cyclically expressed genes. (A)** Additional examples of TFB2 target genes in Vps4 overexpressing cells. A TFB1-dependent example gene, *nurA* (saci_0148) was shown for comparison. **(D)** TFB2 peaks are associated with promoters of upregulated genes in Vps4 overexpression cells. $Log_2$(odd ratios) were calculated for promoters (TSSs) based on whether they are bound by TFB2 and whether genes they control are upregulated. 95% confidence intervals determined by Fisher's method are shown as shaded areas. Increasing numbers of the top TFB2 or TFB1 peaks were included. Data were corrected for the effect of single TFB peaks overlapping with pairs of closely spaced divergent promoters (see methods). **(C)** TFB2 binding at promoters partly overlaps with TFB1.

Venn diagram showing overlap between TFB1 and TFB2 binding at promoters with mapped TSS in Vps4 overexpression cells. TFB peaks overlapping with pairs of closely spaced divergent promoters were counted as single instance. **(D)** Additional examples of TFB2 (orange) and TFB1 (blue) occupancy at TFB2 target genes in the synchronised cells. **(E)** Discriminative motif analysis of BRE-TATA-box motifs occurring promoters with specific binding of TFB2 (n=37) or TFB1 (n=47).

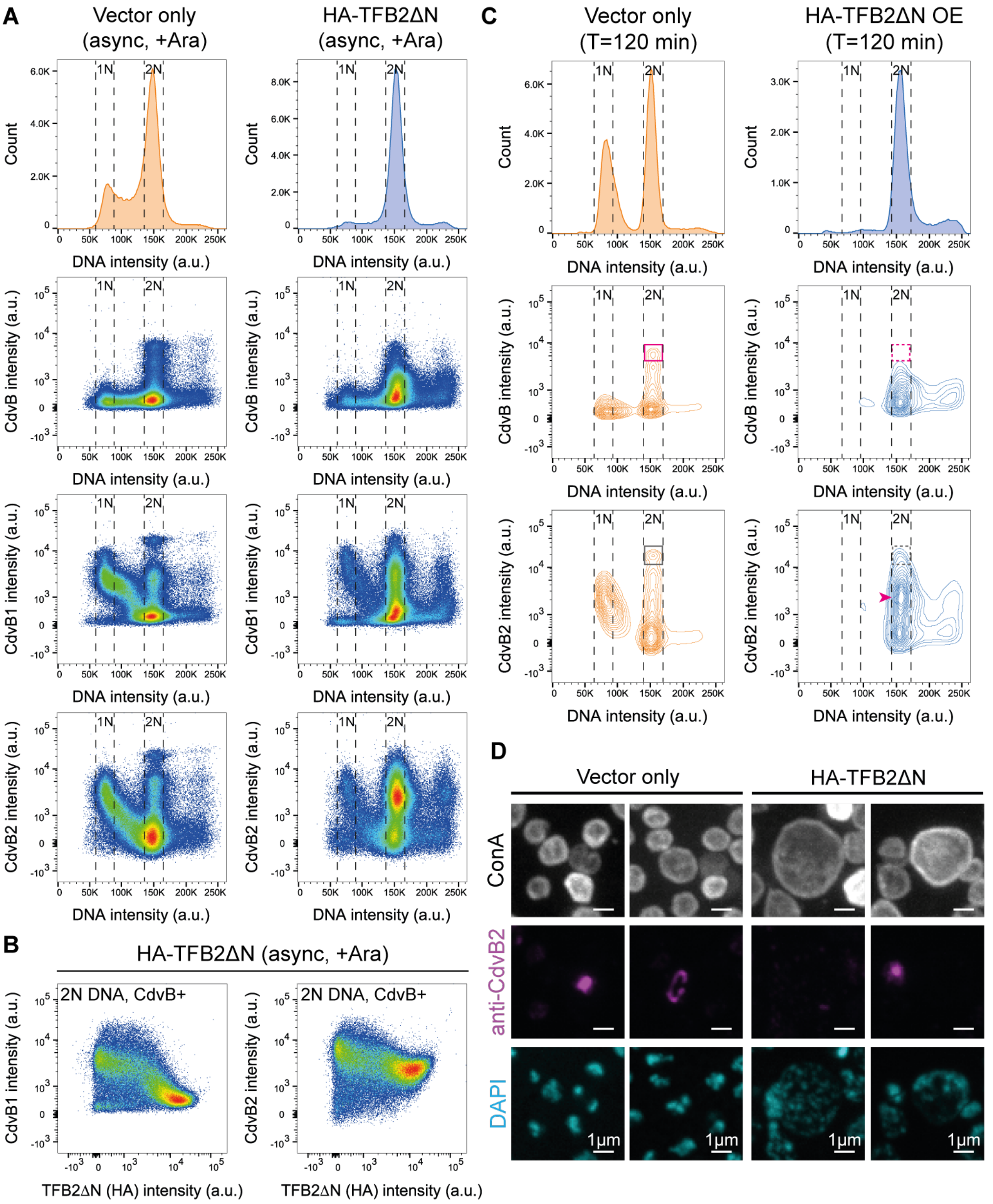


**Supplementary Figure S6. TFB2ΔN mutant disrupts cell cycle progression. (A)(B)** Flow cytometry histograms and scatter plots of vector only control and HA-TFB2ΔN expressing cells. CdvB1 and CdvB2 abundance showed a negative trend with the TFB2ΔN expression level in D-phase (B), indicating that the division gene expression was suppressed by TFB2ΔN mutant. **(C)** Example flow cytometry histograms and contour plots of synchronised cells at around peak of division (arabinose induction from 2 hr before release from acetic acid arrest). The boxes indicated the cells in the pre-constriction phase (red) and constricting phase (grey), which were largely reduced in the TFB2ΔN mutant (right). The arrowhead marks the highest density of cells which contained lower level of CdvB2 than typical late D-phase cells. **(D)** Example microscopy images of immunostained control and TFB2ΔN expressing cells imaged by SoRa spinning disk confocal (maximum projection). TFB2ΔN mutant cells showed clear enlargement in the absence of CdvB2 rings.

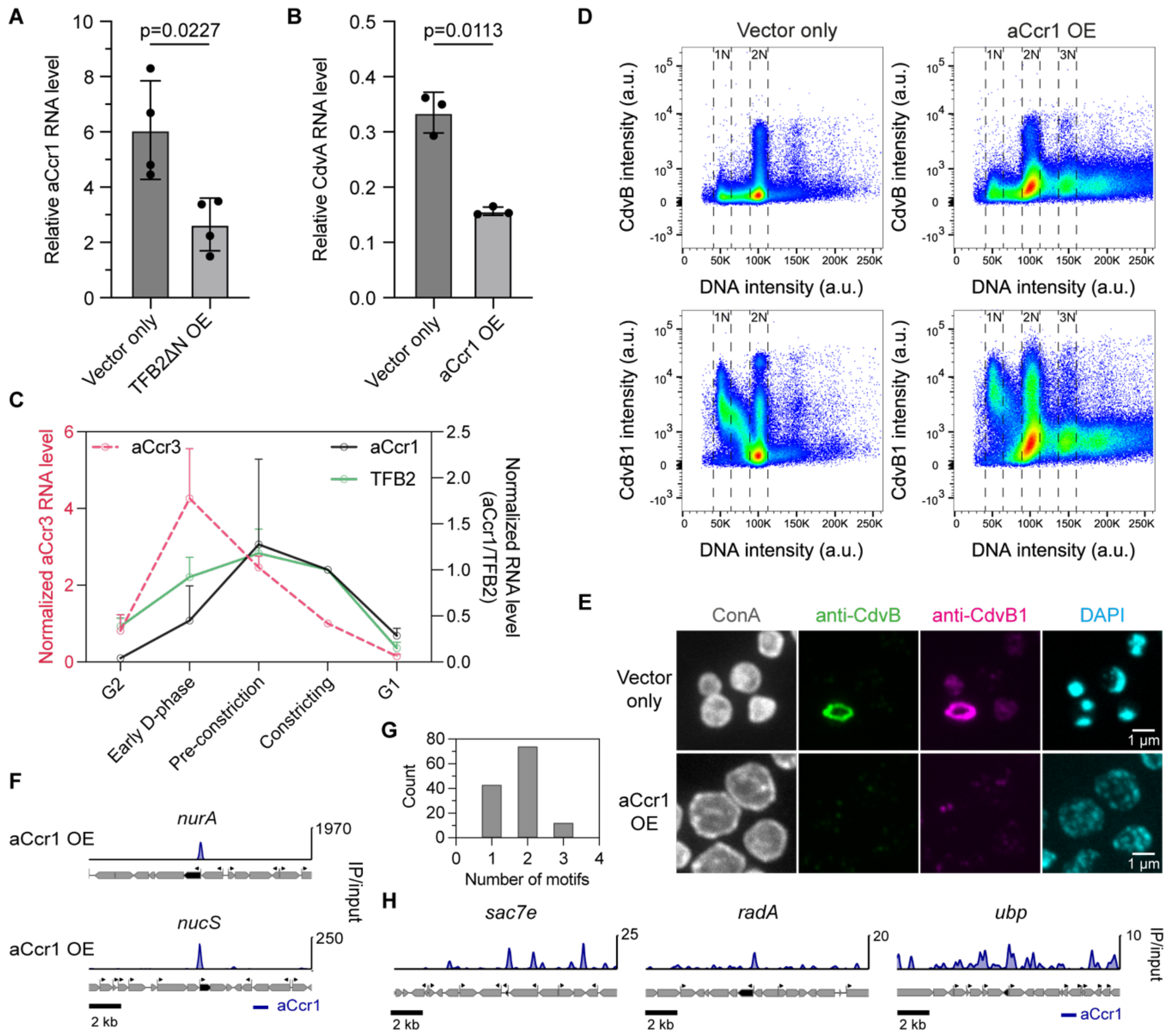


**Supplementary Figure S7. Transcription factor aCcr1 inhibits division gene expression.** **(A)** RT-qPCR analysis of aCcr1 in vector only control and TFB2ΔN expressing mutant (3 hr after arabinose induction). Welch's t-test, N=4 biological replicates. Error bars: mean±SD. **(B)** RT-qPCR analysis of CdvA level in vector only control and aCcr1 OE cells (3 hr after arabinose induction). Welch's t-test, N=3 biological replicates. Error bars: mean±SD. **(C)** Relative RNA level of aCcr1 at different sub-stages during division quantified by RT-qPCR analysis of FACS sorted cells. The transcript level was normalised by the level in the constricting phase. Expression timing of TFB2 and aCcr3 from Fig. 1D and 3C were displayed for comparison. Error bars: mean±SD. N=3 biological replicates. Note that in the aCcr1 curve, the G2 phase data point corresponded to the average of two biological replicates as one replicate showed level below the detection limit. **(D)** Example flow cytometry scatter plots of strains shown in Fig. 4A after 4 hr arabinose induction. Overexpression of aCcr1 represses the expression of CdvB and CdvB1 while DNA re-replication can occur (cells with >2N DNA content). **(E)** Example SoRa spinning disk confocal images of immunostained cells (maximum projection) showing that aCcr1 OE cells inhibit division ring formation accompanied by larger cell size, consistent with failed cell division. **(F)** aCcr1 occupancy at the promoters of the DNA repairing nucleases *nurA* (saci_0148) and *nucS* (saci_0200) in aCcr1 overexpression cells. Note that *nurA* and *nucS* are highly downregulated in the aCcr1 overexpression strain (Fig. 4C) without detectable TFB2 occupancy at their promoter regions in the synchronised dividing cells. **(G)** Number of identified aCcr1 motifs per peaks in aCcr1 OE cells. **(H)** Example aCcr1 occupancy at the promoters of upregulated genes in aCcr1 overexpression condition. These genes were found to be involved in DNA replication, repair or cyclically expressed around S-phase. Note that only 5 upregulated genes ($\log_2(FC) \geq 1$, adjusted p-value <0.01) were found to have aCcr1 binding peaks at the promoter region.

**Supplementary Table S1. Antibodies used in this study.**

| Antibody | Host organism | Dilution | Catalogue number |
|---|---|---|---|
| Anti-CdvB serum | Rabbit | 1:1000 | - |
| Anti-CdvB IgG | Rat | 1:500 | - |
| Anti-CdvB1 IgY | Chicken | 1:1000 | - |
| Anti-CdvB2 IgG (peptide antibody) | Guinea Pig | 1:1000 | - |
| Anti-CdvA IgY | Chicken | 1:1000 | - |
| Anti-saci_0942 (aCcr1) IgG | Rabbit | 1:500-1:1000 | - |
| Anti-StrepII tag Monoclonal Antibody THE™ (clone 5A9F9) | Mouse | 0.2-1 µg/mL | A01732 (GenScript) |
| Anti-HA Monoclonal Antibody (2-2.2.14) | Mouse | 1:1000-1:5000 | 26183 (Invitrogen) |
| Anti-Alba serum | Rabbit | 1:2000 | - |
| Anti-Rabbit IgG, AF488 | Goat | 1:1000 | A11034 (Invitrogen) |
| Anti-Rat IgG, AF488 | Goat | 1:250-1:100 | A-11006 (Invitrogen) |
| Anti-Mouse IgG, AF488 | Goat | 1:200-1:1000 | A11029 (Invitrogen) |
| Anti-Mouse IgG, AF647 | Goat | 1:200-1:1000 | A21235 (Invitrogen) |
| Anti-Chicken IgY, AF546 | Goat | 1:200-1:1000 | A11040 (Invitrogen) |
| Anti-Chicken IgY, AF647 | Goat | 1:1000 | A21449 (Invitrogen) |
| Anti-Guinea Pig IgG, AF546 | Goat | 1:200-1:1000 | A11074 (Invitrogen) |
| Anti-Mouse IgG IRDye 800CW | Goat | 1:10,000 | 926-32210 (LI-COR Biosciences) |
| Anti-Rabbit IgG IRDye 800CW | Goat | 1:10,000 | 926-32211 (LI-COR Biosciences) |

**Supplementary Table S2. Primer pairs for RT-qPCR.**

| Target gene | Forward primer (5' to 3') | Reverse primer (5' to 3') |
|---|---|---|
| SecY (saci_0574) | ACTCTTGCTTGACGAGATGATAC | ACTCTGTACGGAGACTATTCCA |
| 16S rRNA (saci_1300) | CTAGGTGTCGAGTAGGCTTAGAG | CCCGCCAATTCCTTTAAGTTTCA |
| CdvA (saci_1374) | GGACAGATGGAGAAGATAAGGAAG | TGAGCAACTTGGTCCTCTATTG |
| CdvB (saci_1373) | ACTGGTGCATTAAGCGAGAA | TTGGTAACTCTGAAGGTGGATG |
| CdvB1 (saci_0451) | AGACTTCGTTGAAAGGCGTAATG | TCTGTCTTGCCTCTGGTGAATAG |
| CdvB2 (saci_1416) | ATGTGATGAAGGGTGTAATGCCT | TTGCAAAGTCAACTCTAGCTCCA |
| TFB2 (saci_1341) | AGCTTATCACAGGCGGATTTAT | TGCAGGTAATCCCAATCTTTCT |
| aCcr3 (saci_0843) | TCCGGAGCA ATTCCTAGA AGC | ATCACTTACCCATAGTTCCTTCCT |
| aCcr1 (saci_0942) | TGGACAGATATGCAATAAAAACAGG | CGGCACTGTCTCCTTGCTT |
| RNR (saci_1353) | TAGAGGAGAAGGGCGGTTCA | TCCACTATCCCATGCTCCCT |

**Supplementary Table S3. List of plasmids used in this study.**

| Plasmid name | Coding gene | Expression organism | Source |
|---|---|---|---|
| pSVAaraFX-Stop | Vector (no tag) | *S. acidocaldarius* | van der Kolk et al, 2020 |
| pSVAaraFX-HA | Vector (HA-tag) | *S. acidocaldarius* | van der Kolk et al, 2020 |
| pSVAaraFX-saciVps4 | *Saci_1372* | *S. acidocaldarius* | This study |
| pSVAaraFX-saciVps4$^{E209Q}$-His$_6$ | *Saci_1372*$^{E209Q}$ | *S. acidocaldarius* | Hurtig et al. 2023 |
| pSVAaraFX-CdvAΔE3B-HA | *Saci_1374*$^{1-220aa}$ | *S. acidocaldarius* | Parham et al. 2025 |
| pSVAaraFX-HA-LacS-CdvB$^{194-end}$ | *SSO3019, saci_1373*$^{194-end}$ | *S. acidocaldarius* | Kuo et al. 2026 |
| pSVAaraFX-TFB2-FLAG-StrepII | *Saci_1341* | *S. acidocaldarius* | This study |
| pSVAaraFX-TFB2ΔN | *Saci_1341*$^{97-end}$ | *S. acidocaldarius* | This study |
| pSVAaraFX-HA-TFB2ΔN | *Saci_1341*$^{97-end}$ | *S. acidocaldarius* | This study |
| pSVAaraFX-TFB2ΔC-FLAG-StrepII | *Saci_1341*$^{1-192aa}$ | *S. acidocaldarius* | This study |
| pSVAaraFX-aCcr1 | *Saci_0942* | *S. acidocaldarius* | This study |
| pET21a-aCCR1 (Saci_0942) | *Saci_0942* | *E. coli* | This study |

**Supplementary Data S1. Differential gene expression analyses of RNA-Seq experiments.**

**Supplementary Data S2. Summary of peak calling of ChIP-seq experiments.**
